# Abundance, diversity and activity of endogenous retroviruses in the slow loris

**DOI:** 10.64898/2026.06.25.734490

**Authors:** Charles A.G Michie, Hayley Beth Free, Vincent Nijman, Ravinder K. Kanda

## Abstract

Endogenous retroviruses (ERVs) constitute a significant fraction of vertebrate genomes and serve as genomic records of past retroviral infections, while also influencing host biology through regulatory co-option and, in some cases, ongoing retrotransposition. Despite extensive examination of ERVs in haplorrhine primates, equivalent analyses in strepsirrhines remain absent, leaving a substantial gap in our understanding of ERV diversity and evolutionary dynamics across the primate order. Here, we present the first comprehensive characterisation of ERVs in a strepsirrhine primate, identifying 15 Loris Endogenous Retrovirus (LERV) families encompassing 34 subfamilies and over 6,000 insertions in the Nycticebus coucang reference genome. Phylogenetic analyses resolved LERVs into three retroviral genera: betaretroviruses (LERV1–4), type-D betaretroviruses (LERV5–9), and gammaretroviruses (LERV10–15). LERV2a shows multiple hallmarks of recent or potentially ongoing retrotransposition, including a median insertion age of zero, a high proportion of identical LTR pairs, dN/dS ratios comparable to the active retrovirus HTLV, and insertional polymorphism between two conspecific genomes. Comparative genomic screening across Lorisidae revealed that LERV subfamily distribution broadly mirrors estimated insertion ages, with progressively fewer subfamilies detected in more distantly related species. These findings establish a detailed foundation for understanding retroviral evolution in Strepsirrhini and reveal that ongoing retroviral activity is not restricted to haplorrhine primates.

## 1 Introduction

A substantial proportion of vertebrate genomes is derived from viral sequences, the majority of which originate from retroviruses. Retroviruses exist extracellularly with RNA genomes and undergo obligatory reverse transcription into DNA prior to integration into the host genome (Katzourakis & Gifford, 2010). When integration occurs in germline cells, the viral sequence may be inherited by subsequent generations, forming an endogenous retrovirus (ERV; (Lavialle et al., 2013), which is approximately 10kb long containing the internal retroviral genes (*gag, pro, pol, env*) flanked by two identical long terminal repeats (LTRs). Although most ERV insertions are thought to be neutral and lost due to genetic drift, some insertions do become fixed and constitute a significant fraction of the host genome. In mammals ERVs account for approximately 10% of the genome (Lander et al., 2001; McCarthy & McDonald, 2004; Zheng et al., 2024). Some loci are beneficial, and the LTRs have been co-opted as alternative promoters (Chuong et al., 2016; Conley et al., 2008; Fuentes et al., 2018), or retroviral genes have been co-opted and repurposed, as in the example of syncytin gene (Lavialle et al., 2013; Mi et al., 2000). The majority of ERV loci derive from ancient retroviral infections and that lack closely related extant counterparts, however, some ERVs represent recent activity and potentially ongoing exogenous infections (Best et al., 1997; Chu et al., 2023; Herniou et al., 1998; Wang & Han, 2023). ERVs classification is based on their relatedness to extant retroviruses which consists of seven distinct genera: alpha-, beta-, gamma-, delta-, lenti-, spuma-, and epsilon-retroviruses (Gifford et al., 2018). In mammals, most ERVs are derived from and classified as beta- and gamma-retroviruses.

Excluding studies conducted in laboratory mice, the majority of mammalian ERV research has focused on the great apes, including humans (haplorrhine primates). Beyond studies examining associations with disease (Kassiotis & Stoye, 2016, 2025), much of this research has involved investigating the population dynamics and evolutionary consequences of ERVs on the host genome in these primates (Gifford et al., 2005; Jern & Coffin, 2008; Kanda et al., 2013; Kitao et al., 2025; Lee et al., 2014; Y. Li et al., 2022, 2025; Romano et al., 2007). In humans the most recently active ERV family is HERV-K (human endogenous retrovirus) with multiple loci that are human specific (Medstrand & Mager, 1998; Subramanian et al., 2011), several of which are insertionally polymorphic (Subramanian et al., 2011; Wildschutte et al., 2016). At least 30 distinct families of HERVs have been identified (Katzourakis & Tristem, 2005; Mayer et al., 2011). With the exception of HERV-L which inserted prior to the divergence of all mammals (Bénit et al., 1999), and HERV-F/H which inserted prior to the divergence of all primates (Bannert & Kurth, 2006; Izsvák et al., 2025; Patzke et al., 2002), the majority of our understanding of ERVs in mammals comes from the Great Apes, and very little is known across all mammals, or even across the primate order. Examining ERV diversity outside Haplorhini, in the other primate suborder (strepsirrhines – lemurs, galagoes, lorises) will reveal whether the patterns of ERV accumulation documented in these lineages are broadly conserved across primates or are specific to the haplorrhine clade.

Few studies have investigated ERVs specifically in Strepsirrhini. In the Ring tailed lemur (*Lemur catta*), approximately 19% of transposable elements were determined to be ERVs (Palmada-Flores et al., 2022). In mouse lemurs (*Microcebus* spp.) three gammaretrovirus and betaretrovirus families were identified with average ages of insertion ranging from 44 to 60 million years (Kessler et al., 2023). In Coquerel’s sifaka (*Propithecus coquereli*) a betaretrovirus named “Prosimian retrovirus 1” was identified and aged to 4.5 million years ago and is represented by only three genomic copies (Apakupakul et al., 2021). Endogenous lentiviruses have been identified in the Fat-tailed dwarf lemur (*Cheirogaleus medius*) and the Gray mouse lemur (*Microcebus murinus*) and lie basal to simian immunodeficiency virus (SIV) and human immunodeficiency virus (HIV; (Gilbert et al., 2009). The few reported ERVs in Strepsirrhini all appear ancient with no evidence of recent or ongoing insertions, which is unexpected given the relatively recent ERV activity observed in the great apes (Holloway et al., 2019; Subramanian et al., 2011). A comprehensive examination of ERVs in a Strepsirrhini species is necessary to ascertain whether our understanding of ERV dynamics from observations in Haplorhini are consistent.

Slow lorises (*Nycticebus* spp.) are nocturnal venomous Strepsirrhini primates native to southeast Asia. Slow loris taxonomy is complicated: the species are largely cryptic, with minor morphological differences that are difficult to observe. (Nekaris & Starr, 2015), and the phylogenetic relationships do not consistently align with geographic or morphological boundaries (Blair et al., 2023; Pozzi et al., 2015). It is further complicated by the historical changes in taxonomy, where until 1971 all slow lorises were classified as *N. coucang* with multiple subspecies (Groves, 1971) and were not elevated to species status until 2001, including *N. bengalensis* (Groves, 2001). This has led to some mislabelling as it is believed that the reference *N. coucang* genome on NCBI (GCA_027406575.1) is actually a *N. bengalensis* (Michie et al., 2025). There are currently 10 recognised slow loris species, and all are classified as globally threatened (Vulnerable, Endangered, or Critically Endangered) on the IUCN Red List, primarily due to habitat loss and the illegal wildlife trade (Moore et al., 2014; Nekaris & Burrows, 2020). In addition, through their involvement in the illegal pet trade (Nekaris et al., 2010) and their use in traditional medicine (Starr et al., 2010; Thạch et al., 2018), slow lorises experience frequent contact with humans and other animals, raising the possibility of exposure to, or transmission of, infectious agents including viruses. Coupled with their endangered status, there is interest in genomic and population level analysis to better inform conservation efforts, which provides a timely opportunity to examine their ERV content.

There has been no comprehensive study of ERVs within slow lorises specifically, and they have only been included as a species in a large dataset where their specific ERVs have not been examined in any detail (Gifford et al., 2005; Kitao et al., 2025; Y. Li et al., 2022, 2025).; only two of these studies included any information on the ERV sequences identified. Gifford et al., (2005) included a partial amino acid sequence of the reverse transcriptase found in *N. coucang* labelled as “RV Slow Loris” and Li et al., (2022) included the genomic locations of an “Unknown HERV” and a HERV-HF found in *N. bengalensis* both of which did not have all major genes identified. Michie et al., (2025) identified the first complete full length ERV (LERV1) in *Nycticebus* which was a betaretrovirus which is most closely related to HERV-K. Comparison between the reference *N. coucang* (GCA_027406575.1) and *N. bengalensis* (GCA_023898255.1) genomes identified high similarity among LERV1 loci which was unexpected given their ∼5 MYA divergence time and ages of LERV1 insertions. Combined with mitochondrial DNA evidence, it was concluded that the current reference *N. coucang* genome is most likely a *N. bengalensis* (Michie et al., 2025).

Analysis of ERVs in slow lorises enables direct comparison of retroviral evolution across the deep Haplorhini–Strepsirrhini split, offering insight into how host lineage divergence has shaped ERV diversity and persistence in primates. Examining ERVs in slow lorises allows assessment of whether retroviral activity in strepsirrhines is predominantly ancient or whether more recent, potentially active insertions persist in these understudied primates.

## 2 Results

### 2.1 Identification and Classification of LERV Families in *N. coucang*

We ran RepeatModeler2 (RM2) (Flynn et al., 2020) on the reference *N. coucang* genome (GCA_027406575.1). From the LTR structural detection round this produced a total of 45 Type=LTR and 71 Type=INT families as possible ERV family candidates. If a Type=LTR was found to have a correspond Type=INT based on genomic co-ordinates, the Type=LTR was kept and used for downstream analysis. The families identified by RM2 were grouped into three categories:

Type=LTR with a corresponding Type=INT (n=35)
Type=INT with no corresponding Type=LTR (n=39)
Type=LTR with no corresponding Type=INT (n=9),

Not all RM2 identified families that fitted neatly into these categories – in some instances several Type=INT families corresponded to one Type=LTR family, or the reverse (one Type=INT corresponding to several Type=LTR families).

BLASTn results of the candidate ERV LTRs and internal regions confirmed identification of potential ERV families for 33/35 RM2 families in category 1, 18/39 in category 2, and 9/9 from category 3 after removal of redundancy arising from overlapping BLASTn hits. BLASTn hits had an extra 14,000 bases of flanking region extracted on either side to find complete full-length (CFL) ERVs containing all major genes. Conserved domains in each locus were identified with rpstBLASTn using the NCBI conserved domain database. CFL ERV loci were determined using the criteria of conserved domains present in each of the four genes, and two complete LTRs flanking the internal region – this left 30 candidate ERV families. Within these 30 potential families there was a high degree of similarity between some families, and final classification of families and subfamilies was based on the identity of sequences determined by CD-hit (Fu et al., 2012; W. Li & Godzik, 2006) and manual examination. As recombination is a possibility each gene was examined individually. If at least one of *gag*, *pol* or *env* shared more than 80% nucleotide identity, elements were assigned to the same family., whereas if the entire ERV (LTR, *gag*, *pro*, *pol*, or *env*) was above 80% nucleotide identity they were the same subfamily. This left a total of 34 subfamilies across 15 families of ERVs, here named LERV (Loris Endogenous Retrovirus) which included the previously identified LERV1 family (Michie et al., 2025) (Table 1). Subfamilies were most commonly differentiated based on LTR, where the internal genes were still over 80% similar but the LTRs were not.

**Table 1:** Correspondence between RepeatModeler2-identified repeat families and curated LERV family and subfamily designations. For each RM2 family, the assigned LERV family, resulting subfamilies, and the genomic regions used to discriminate subfamilies are shown. Where a dash (–) appears in the subfamily column, no subfamilies were identified within that family.

| Final LERV family | Final LERV subfamilies | Subfamily classification based on |
| --- | --- | --- |
| 1 | LERV1a, LERV1b | LTR |
| 2 | LERV2a, LERV2b, LERV2c, LERV2d, LERV2e | LTR |
| 3 | LERV3a, LERV3b | LTR |
| 4 | LERV4a, LERV4b, LERV4c, LERV4d | LTR, gag |
| 5 | - | - |
| 6 | - | - |
| 7 | - | - |
| 8 | LERV8a, LERV8b, LERV8c | LTR, pol |
| 9 | LERV9a, LERV9b | LTR, gag, env |
| 10 | - | - |
| 11 | - | - |
| 12 | - | - |
| 13 | LERV13a, LERV13b, LERV13c, LERV13d, LERV13e, LERV13f | LTR, gag, pol |
| 14 | LERV14a, LERV14b | LTR, pol |
| 15 | LERV15a, LERV15b | LTR |

There was substantial variation in the number of loci among LERV families and subfamilies, with some having fewer than 10 insertions and others having over a thousand (Table 2). For every LERV family, target site duplications (TSDs) could be identified for the majority of the loci. The length of the TSD varied between families and ranged from four to six base pairs. The age of insertion for ERVs can be estimated from the divergence of LTRs as they are identical at the time of insertion. The median age of insertion for each LERV family ranged from 0 (LERV2a and LERV2b) to 39.26 million years old (LERV12) (Figure 1). When considering only LERV subfamilies with more than 10 CFL and solo LTRs loci, a moderate negative linear correlation was observed between percentage of insertions with matching TSDs and the median estimated age of insertion (R^2^ = 0.488, p <0.0001). The percentage of solo LTRs to CFL loci varied from 25% to 98.77% and there was no strong linear correlation with median estimated age of insertion (R^2^ = 0.018, p = 0.45). There was no linear correlation between age and the number of loci in a family (R^2^ = 7×10^−5^, p = 0.96).

**Figure 1.**
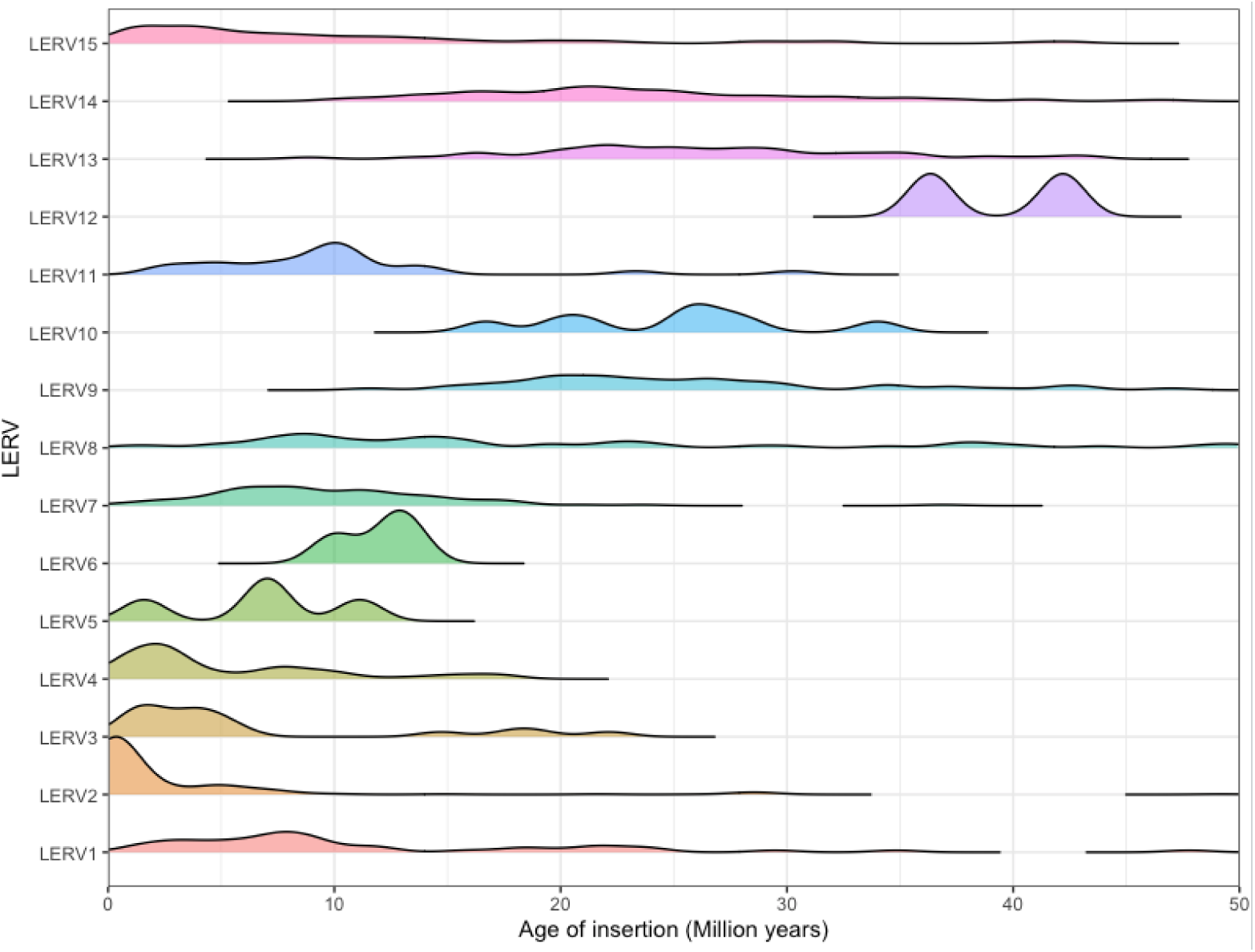
Estimated insertion age distributions of LERV subfamilies in the N. coucang reference genome. Insertion ages were estimated from LTR sequence divergence of complete full-length (CFL) elements, using a neutral substitution rate of 1.72 × 10 substitutions/site/year (Campbell et al. 2021).

**Table 2:** Copy number, target site duplication (TSD) matching rate, and estimated insertion age of LERV subfamilies identified in the N. coucang reference genome. The number of solo LTR and complete full-length (CFL) insertions is shown for each subfamily, with the percentage of loci exhibiting matching TSDs in parentheses. Insertion ages were estimated from LTR sequence divergence using a neutral substitution rate of 1.72 × 10 substitutions/site/year (Campbell et al. 2021) and are expressed in millions of years ago (MYA); median values are shown with the 75% interquartile range in brackets.

| LERV subfamily | Number of Solo (% with matching TSD) | Number of CFL (% with matching TSD) | Median age of CFL in MYA with matching TSDs (75% IQR) | Retroviral genera |
| --- | --- | --- | --- | --- |
| LERV1a | 1127 (74) | 14 (71) | 19 (13.63-21.8) | Betaretrovirus |
| LERV1b | 411 (87) | 32 (91) | 6.98 (3.55-9.04) | Betaretrovirus |
| LERV2a | 60 (97) | 159 (97) | 0 (0-0.29) | Betaretrovirus |
| LERV2b | 89 (97) | 20 (100) | 0 (0-0.8) | Betaretrovirus |
| LERV2c | 174 (95) | 21 (90) | 0.83 (0.56-1.36) | Betaretrovirus |
| LERV2d | 97(97) | 20 (100) | 1.15 (0.87-1.39) | Betaretrovirus |
| LERV2e | 5(80) | 2(100) | 1.92(1.47-2.36) | Betaretrovirus |
| LERV3a | 115 (94) | 4 (100) | 1 (0.43-1.57) | Betaretrovirus |
| LERV3b | 118 (92) | 9 (100) | 1.77 (1.18-3.58) | Betaretrovirus |
| LERV4a | 22 (100) | 4 (100) | 3.14 (2.47-3.95) | Betaretrovirus |
| LERV4b | 83 (95) | 4 (75) | 2.57 (1.67-3.57) | Betaretrovirus |
| LERV4c | 77 (100) | 14 (100) | 2.62 (2.05-4.68) | Betaretrovirus |
| LERV4d | 34 (97) | 9 (100) | 1.79 (0.94-2.69) | Betaretrovirus |
| LERV5 | 20 (95) | 4 (100) | 7.03 (5.59-8.14) | Type-D Betaretrovirus |
| LERV6 | 1 (100) | 3 (100) | 12.49 (11.24-12.91) | Type-D Betaretrovirus |
| LERV7 | 149 (94) | 45 (91) | 9.63 (5.24-14.01) | Type-D<br>Betaretrovirus |
| LERV8a | 95 (92) | 33 (82) | 16.06 (10.08-35.35) | Type-D<br>Betaretrovirus |
| LERV8b | 21(100) | 9 (89) | 16.98 (11.96-19.84) | Type-D<br>Betaretrovirus |
| LERV8c | 38 (84) | 10 (100) | 9.02 (7.2-20.42) | Type-D<br>Betaretrovirus |
| LERV9a | 108 (86) | 30 (97) | 24.52 (19.48-30.37) | Type-D<br>Betaretrovirus |
| LERV9b | 59 (81) | 26 (88) | 23.69 (20.48-28.35) | Type-D<br>Betaretrovirus |
| LERV10 | 93 (75) | 4 (100) | 23.42 (20.05-26.63) | Gammaretrovirus |
| LERV11 | 9 (89) | 13 (100) | 7.46 (4.26-11.61) | Gammaretrovirus |
| LERV12 | 25 (68) | 2 (100) | 39.26 (37.78-40.72) | Gammaretrovirus |
| LERV13a | 20 (80) | 1 (100) | 27.32 | Gammaretrovirus |
| LERV13b | 20 (90) | 10 (100) | 29.94 (22.18-35.5) | Gammaretrovirus |
| LERV13c | 9 (78) | 4 (100) | 36.37 (28.3-49.59) | Gammaretrovirus |
| LERV13d | 63 (81) | 9 (78) | 26.94 (22.38-34.58) | Gammaretrovirus |
| LERV13e | 337 (84) | 6 (100) | 25.29 (24.83-31.57) | Gammaretrovirus |
| LERV13f | 136 (79) | 28 (93) | 26.23 (21.04-31.37) | Gammaretrovirus |
| LERV14a | 153 (79) | 11 (73) | 32.84 (23.8-34) | Gammaretrovirus |
| LERV14b | 137 (82) | 47 (83) | 31.71 (26.1-42.95) | Gammaretrovirus |
| LERV15a | 338 (88) | 64 (98) | 10.74 (7.06-14.55) | Gammaretrovirus |
| LERV15b | 272 (91) | 97 (96) | 4.17 (1.91-11.12) | Gammaretrovirus |

**Table 3.** Nonsynonymous-to-synonymous substitution rate ratios (dN/dS) for internal retroviral genes estimated using the one-ratio (M0) model in CODEML. a) dN/dS values for LERV families in which full length open reading frames could be reconstructed. Published dN/dS values for HERV-K (HML2) are from Belshaw et al. (2004); Human T-lymphotropic virus (HTLV) values are shown as a reference for an extant, actively replicating retrovirus. N/A indicates genes for which reliable dN/dS estimation was not possible due to insufficient data or alignment ambiguity. b) Separate dN/dS values for the subfamilies of LERV2. LERV2a active candidates are defined as insertions with identical LTRs, matching TSDs, and complete open reading frames requiring no manual correction.

| Retroviral Family | Number of elements | Median age (MYA) | *gag* | *pro* | *pol* | *env* |
| --- | --- | --- | --- | --- | --- | --- |
| LERV1 | 40 | 8.20 | 0.736 | 0.493 | 0.526 | 0.553 |
| LERV2 (all) | 217 | 0 | 0.185 | 0.074 | 0.159 | 0.213 |
| LERV3 (all) | 13 | 1.48 | 0.288 | 0.143 | 0.169 | 0.194 |
| LERV4 (all) | 31 | 7.03 | 0.300 | 0.202 | 0.231 | 0.316 |
| LERV7 | 42 | 9.18 | 0.929 | N/A | 0.606 | 0.564 |
| LERV8 (all) | 52 | 21.97 | 0.723 | 0.781 | 0.740 | 0.742 |
| LERV15 (all) | 155 | 7.63 | 0.958 | N/A | 0.846 | 0.776 |
| HERV-K (HML2) | 44 | N/A | 0.53 | N/A | 0.47 | 0.62 |
| Human T-lymphotropic virus | 49 | 0 | 0.105 | N/A | 0.135 | 0.148 |

b)
| LERV2 subfamily | Number of elements | Median age (MYA) | *gag* | *pro* | *pol* | *env* |
| --- | --- | --- | --- | --- | --- | --- |
| LERV2a (all) | 158 | 0 | 0.169 | 0.083 | 0.134 | 0.188 |
| LERV2a active candidates | 31 | 0 | 0.155 | 0.121 | 0.134 | 0.148 |
| LERV2b | 18 | 0 | 0.155 | 0.076 | 0.122 | 0.279 |
| LERV2c | 20 | 0.83 | 0.206 | 0.069 | 0.203 | 0.273 |
| LERV2d | 19 | 1.15 | 0.159 | 0.067 | 0.202 | 0.138 |

### 2.2 Phylogenetic placement of LERV families

The phylogenetic placement of LERVs was based on amino acid alignments with conserved domains from *pol* (reverse transcriptase) and *env* (transmembrane subunit) genes followed by phylogenetic tree construction with MrBayes (Huelsenbeck & Ronquist, 2001). This indicated our identified ERVs represented betaretroviruses, type-D betaretroviruses and gammaretroviruses (Supplementary Figure 1; Supplementary Table 1; Table 2). After these initial phylogenetic trees to determine genera placement, phylogenetic trees were also constructed for the full-length nucleotide *pol* and *env* genes to provide a more detailed insight into the relationship with other retroviruses. Amongst the betaretroviruses LERV1 clustered most closely with HERV-K families as found in Michie et al. (2025), while LERV2, LERV3 and LERV4 formed a distinct clade most closely related to rodent betaretroviruses (Figure 2).

**Figure 2:**
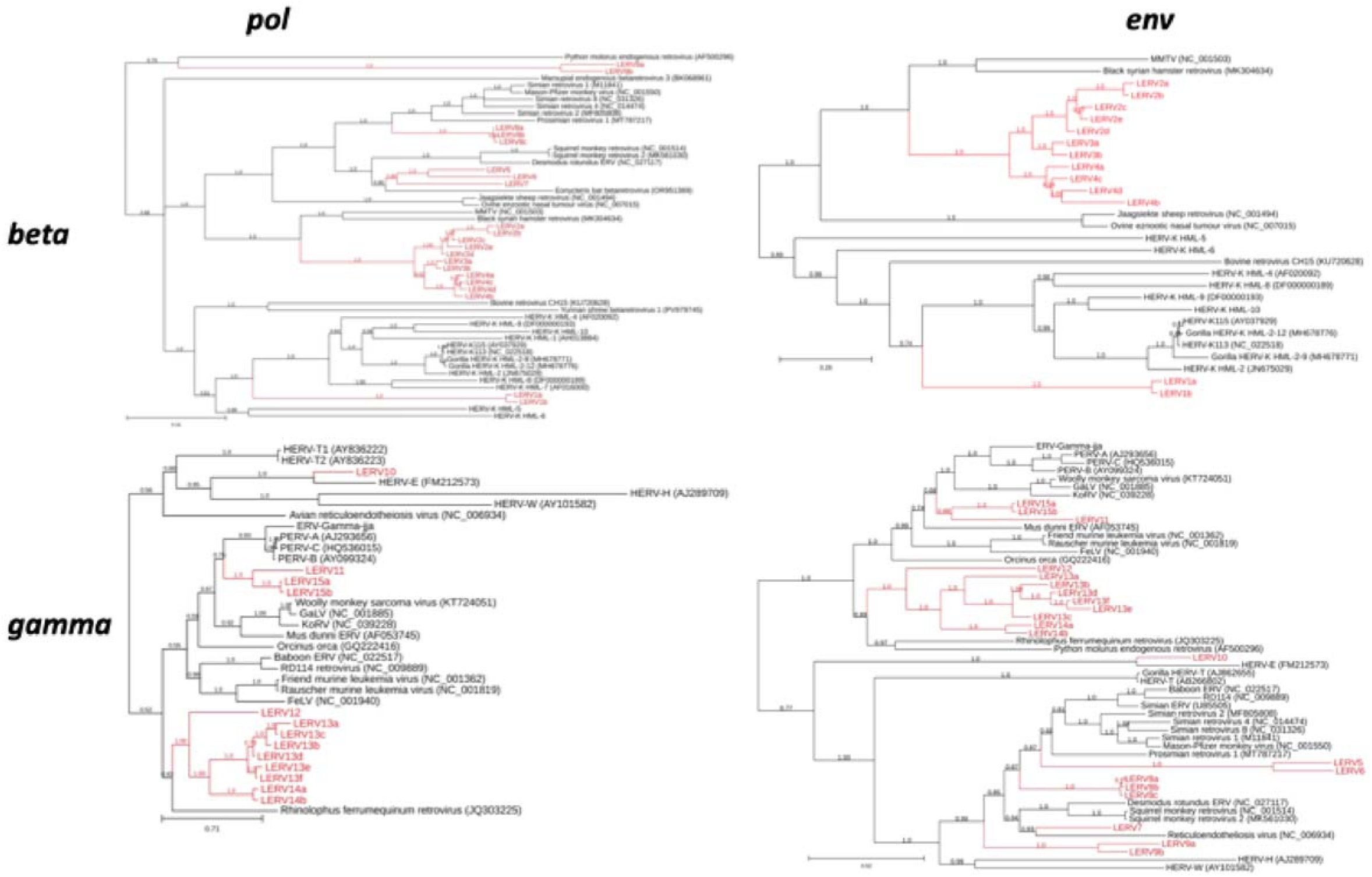
Bayesian phylogenetic tree of LERV families inferred from pol and env whole nucleotide region rooted using Avian Leukosis Virus as an outgroup (not shown for visual purposes). LERV families identified in this study are highlighted in red. Reference retroviral and endogenous retroviral sequences are labelled by common name and GenBank accession number. Phylogenetic trees inferred from gag and pro sequences are provided in Supplementary Figure 2.

Amongst the Type-D Betaretroviruses (beta-type *pol*, gamma-type *env*), phylogenetic analyses revealed overall relationship with other primate type-D betaretroviruses, however these relationships varied between genes. While the gag and pol phylogenies were in agreement, the env phylogenies showed substantial phylogenetic incongruence relative to other endogenous retroviruses. For example, in the *pol* gene, LERV5, LERV6 and LERV7 form a clade most closely related to Squirrel monkey retrovirus, whereas in *env,* only LERV7 retains this relationship with LERV5 and LERV6 instead clustering more closely with simian retroviruses. LERV8 was consistently most closely related to simian retroviruses in all genes. LERV9 also exhibited gene-dependent phylogenetic placement, clustering most closely with *Python molorus* ERV in *pol*, while appearing basal to all primate Type-D betaretroviruses in *env*. Additionally LERV5 may correspond to the previously identified “RV Slow Loris” (Gifford et al., 2005), which was originally reported based on a short amino acid sequence (165aa – 95.15% identity with LERV5) within the reverse transcriptase region (Supplementary Figure 1). RV Slow Loris was originally classified as a betaretrovirus; additional analysis of the LERV5 *env,* indicates this is a type-D betaretrovirus. Within the gammaretroviruses, LERV10 clustered most closely with HERV-E across all genes. LERV11 and LERV15 form a clade across all genes; however, their phylogenetic relationships to other retroviruses varied by gene. In *pol* this clade was most closely related to porcine endogenous retroviruses (PERVs), whereas in *env,* PERVs clustered more closely with koala retrovirus (KoRV) and gibbon ape leukemia virus (GaLV) with LERV11 and LERV15 having a more basal position to this clade. LERV12, LERV13 and LERV14 form a clade in each gene and in *pol* and *env* were most closely related to *Rhinolophus ferrumequinum* retrovirus.

### 2.3 LERVs in related species

To detect the presence of the novel LERVs identified here outside of the Nycticebus spp., the consensus sequence of each LERV subfamily was analysed using BLASTn with a cut-off of 90% query coverage and 80% sequence similarity (including the LTR) against publicly available reference genomes of other primate species (Figure 3). These strict parameters were used to confirm that BLAST hits were of the same LERV family. To assess insertionally polymorphic loci in the potentially active LERV2a subfamily, LTRs from complete full-length insertions with 1000bp of flanking region were used to query genomes from conspecific and closely related species using BLAST to find potential pre-integration sites. In the *Nycticebus bengalensis* genome (NCBI Accession number - GCA_023898255.1), shared LERV2a insertions and corresponding pre-integration sites were identified, indicating insertional polymorphism between the two genomes. LERV2a was not found in any other genomes. With all LERV families, in other members of the subfamily Lorisinae, 29 of the 34 LERV subfamilies were found in *Xanthonycticebus pygmaeus* (∼10 MYA divergence; NGDC GWH Accession number - GWHBCHX00000000), while only 15 were found in *Loris tardigradus* (∼29 MYA divergence; NCBI Accession number - GCA_023783135.1). Notably, for all LERV8 subfamilies in *L. tardigradus*, the *gag* and *pol* genes showed over 80% similarity, but the *env* gene showed only 50% similarity, suggesting that recombination may have occurred. In more distantly related taxa, only three LERV subfamilies were found in *Perodictus potto* (∼38 MYA divergence; NCBI Accession number - GCA_963574655.1) within the family Lorisidae. Within the superfamily Lorisoidea, *Otolemur garnettii* (∼40 MYA divergence; NCBI Accession number - GCA_000181295.3) contained only one LERV subfamily (LERV10). Finally, within the suborder Strepsirrhini, *Lemur catta* (∼59 MYA divergence; NCBI Accession number - GCA_020740605.1) showed no detectable LERVs. The estimated insertion dates broadly correspond to the species in which each LERV was detected, the largest exception is LERV10 whose oldest insertion is ∼27 MYA, 13 MYA after the divergence with *O. garnettii*. LERV10 only had four CFL identified, therefore it is plausible these were only the most intact and therefore the youngest elements that were found.

**Figure 3:**
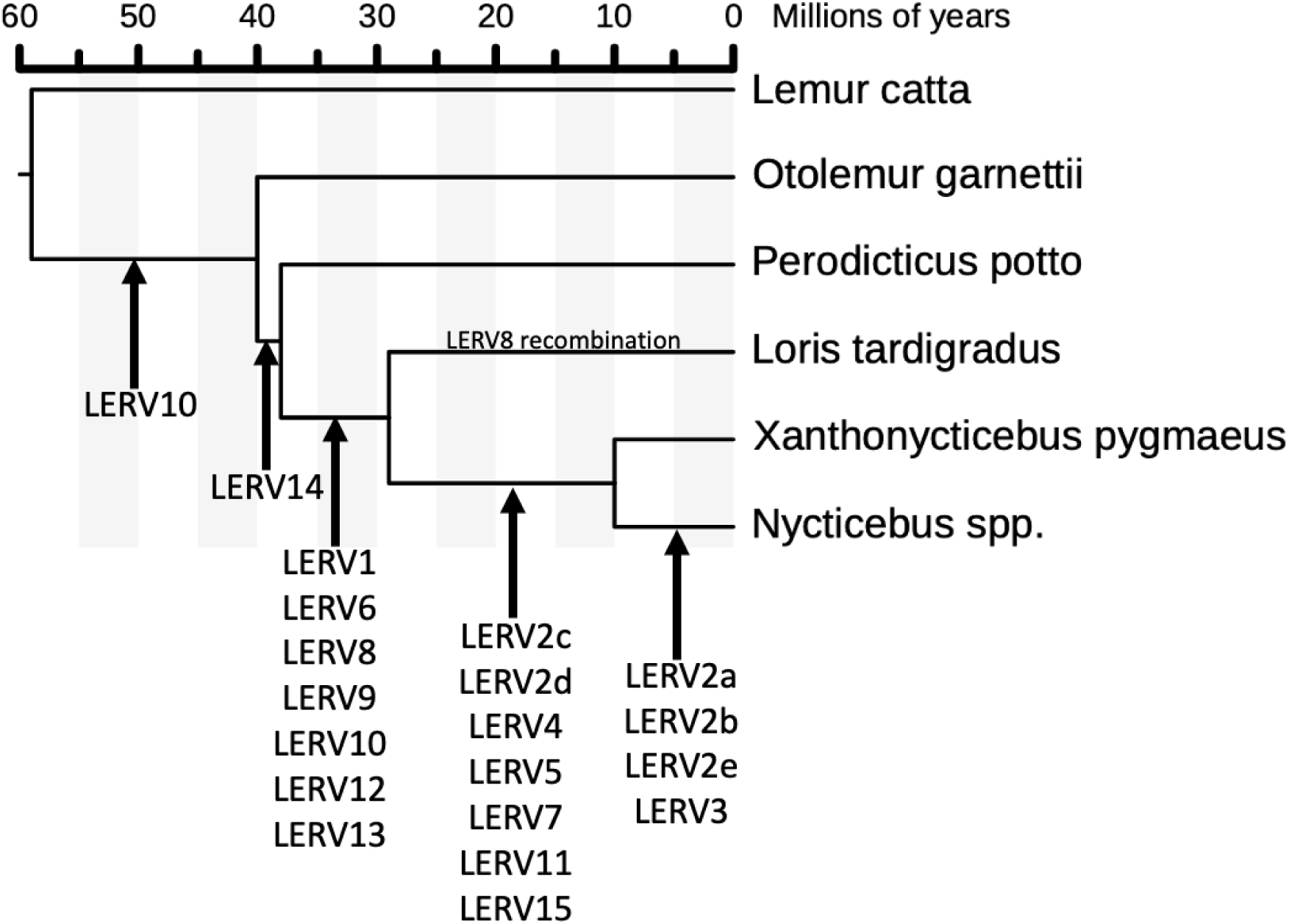
Presence and absence of LERV subfamilies across Lorisidae and related strepsirrhine taxa. Arrows indicate detection of a given LERV subfamily in the queried genome (≥80% sequence identity, ≥90% query coverage).

### 2.4 Related viruses to LERVs in non-primate mammals

Phylogenetic analyses using published sequences can help assign novel retroviruses to a genus but may not identify their closest relatives due to limited ERV sampling across species. To detect closely related viruses, consensus sequences from LERVs unique to *Xanthonycticebus* and *Nycticebus* were queried against the mammalian NCBI RefSeq genome database using BLASTn. BLASTn searches identified rodent-associated sequences as the closest matches for LERV2–4, with *Jaculus jaculus* consistently representing the top hit (>70% query coverage, >65% identity). However, full-length comparisons indicated moderate overall similarity (63.1%), exceeding that observed between LERV2-4, with reference and previously published retroviruses such as MMTV (Figure 4). In contrast, LERV5 showed broader affinities, with top hits in both rodents (*Arvicanthis niloticus*) and Eulipotyphla (*Erinaceus europaeus*), exhibiting 56.7% and 66.6% full-length identity, respectively. Sliding window analysis revealed that similarity to rodent sequences was concentrated in the pro and pol genes. LERV7 displayed highest similarity to sequences identified in bats (*Pipistrellus kuhlii*) and the pangolin (*Manis javanica*), with 69.7% and 66.6% identity across the provirus. For LERV11 and LERV15, the closest matches were again found in bat (*Myotis myotis*) and rodent (*Sciurus vulgaris*) genomes; these sequences showed higher similarity to LERV15 (79.3% and 77.1% identity) than LERV11 did despite the two being each other’s closest related ERV. Sliding-window similarity profiles show noticeable decreases in similarity near the receptor binding domain in *env* and in *gag* between the matrix and capsid domains.

**Figure 4:**
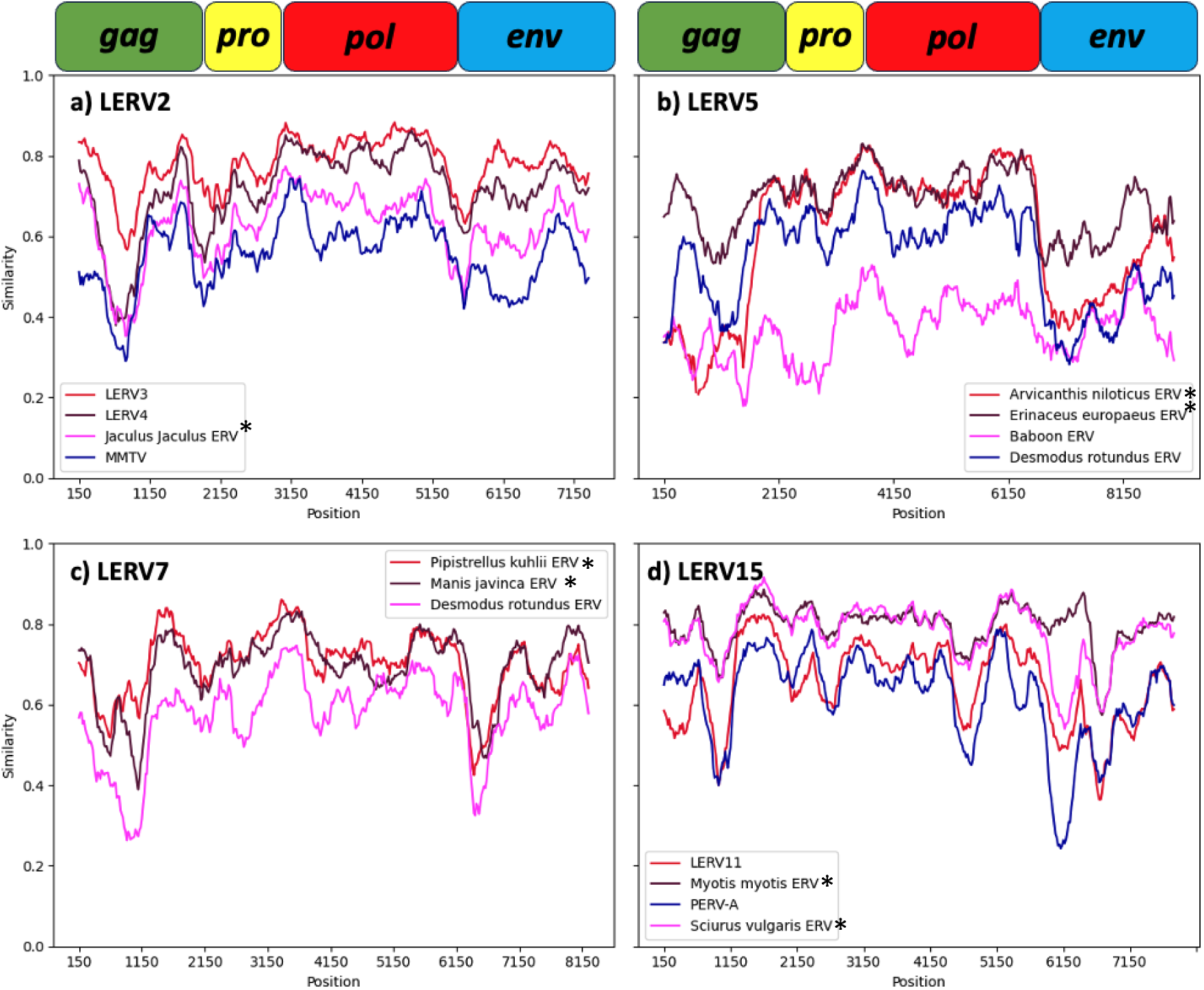
Sliding-window nucleotide similarity between selected LERV families and their closest genomic matches identified in other mammalian species. Similarity was calculated using the Hamming distance model with a 200 bp sliding window and 20 bp step size, implemented in SimPlot++ (Samson et al. 2022). The approximate positions of retroviral genes (gag, pro, pol, env) are indicated above each panel. ERVs found through BLASTn of NCBI genome database are annotated with *.

**Figure 5:**
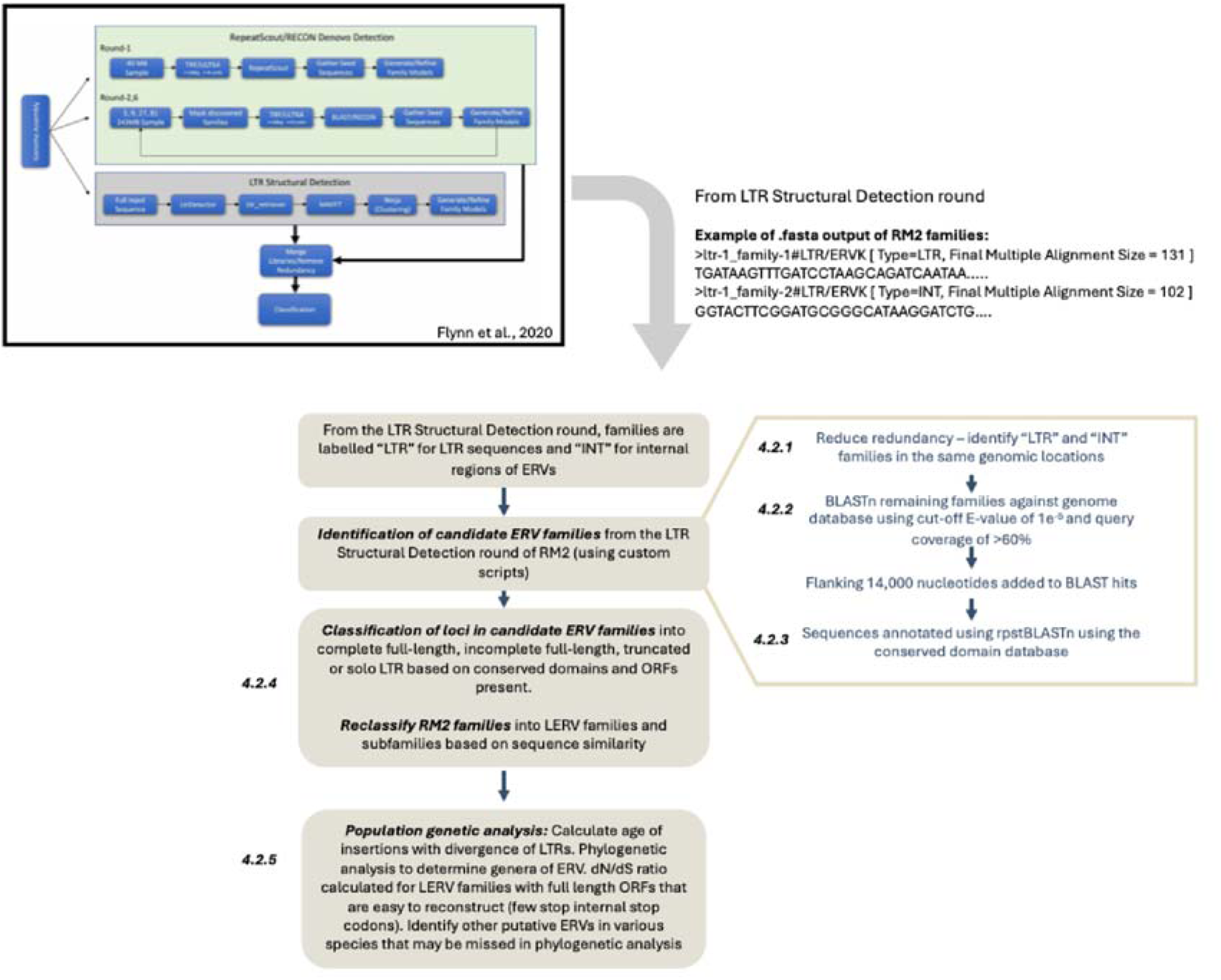
Overview of methods used for the identification of novel endogenous retroviruses in the slow loris. The output from the LTR Structural detection round from RepeatModeler2 (Flynn et al., 2020) is passed through various subsequent curation steps and processed for further analysis.

### 2.5 dN/dS ratios of LERV families

The dN/dS ratio can provide insight into the selective pressures acting on ERV genes within the host genome, where stronger purifying selection is consistent with more recent activity. The dN/dS ratio was calculated with CODEML (Yang, 2007) on families whose entire open reading frames (ORFs) for each gene could be identified. Separate analysis was performed for active candidates of LERV2a, defined as insertions with identical LTRs, matching TSDs and complete ORFs which did not need editing (Table 4). As branch lengths of phylogenies were often zero, branch specific models were not applicable and only the one-ratio model could be completed providing the dN/dS for the whole tree and sequences included. Resulting dN/dS values were compared with published values for HERV-K HML2 (Belshaw et al., 2004) and with those from an extant active retrovirus Human T- lymphotropic virus (HTLV) (Supplementary Table 2).

**Table 4.** Proportion of codon sites under statistically significant purifying selection (p < 0.05) estimated using the Fixed Effects Likelihood (FEL) method (Murrell et al. 2012). Values represent the percentage of codons within each gene showing evidence of purifying selection across all included sequences.

| Retroviral (sub)Family | *gag* | *pol* | *env* |
| --- | --- | --- | --- |
| LERV1 | 8.18% | 9.89% | 18.57% |
| LERV2a active candidates | 27.18% | 19.02% | 29.1% |
| Human T-lymphotropic virus | 14.29% | 13.62% | 11.7% |

Across all genes, a moderately strong positive correlation was observed between median age of insertion and all PAML-derived dN/dS values (R^2^=0.611, p < 0.0001). Among individual genes, *env* exhibited the strongest correlation with age (R^2^=0.7059, p = 0.0003; *pro* excluded due to several N/A values). Several LERV families exhibited marked differences in dN/dS ratios between the *env* and *pol* genes; for example, in LERV2b, dN/dS values were higher in *env* than in *pol*, and in LERV2d, dN/dS values were lower in *env* relative to *pol*. The dN/dS values of the putatively active LERV2a candidates were comparable to those observed for the active HTLV, especially in *pol* and *env*. When separating *gag* into the matrix and capsid domain, the matrix domain had a much lower (0.099) than the capsid domain (0.128). Using FEL (Murrell et al., 2012) the active candidates of LERV2a had a higher proportion of variant codon sites under purifying selection (p<0.05) across all comparable genes (Table 5).

### 2.6 Distribution across chromosomes

Across every LERV family there was unequal distribution across chromosomes (Supplementary Table 3). Chromosomes 20 and X was enriched for LERV insertions where they had 2.34x and 1.47x (BH adjusted p-value < 0.0001) respectively more LERV insertions that would have been expected given the chromosome size. Chromosome 20 had the third most LERV insertions of any chromosome despite being the 21^st^ largest. Chromosomes 2,3,4,7,8,9,11,12,16,17,18,19,21,22 and 23 had significantly less LERV insertions than what would be expected by chance (all BH adjusted p-value < 0.05). There was not a significant difference in the number of insertions for any orientation across the whole genome or on any individual chromosome (all BH adjusted p-value > 0.15).

## 3. Discussion

This study provides the first detailed examination of the diversity and abundance of ERVs in a strepsirrhine primate. Using RM2, followed by in-depth examination of elements identified in the reference *N. coucang* genome, over 6000 ERV insertions were found, forming 15 families and 34 sub-families. This is substantially fewer than the ∼700000 repeats in humans from 30 to 50 families (https://herv.img.cas.cz/statistics; Mayer et al., 2011), and may reflect bias in identifying more recently inserted families containing fewer mutations, thus rendering them easier to identify and accurately determine to be a true ERV. It may also be because a large majority of predicted HERVs are not true ERVs as only the *pol* gene is often found (Y. Li et al., 2025). The pattern observed in humans, where most ancient insertions are gammaretroviruses and more recent insertions are betaretroviruses (Chabukswar et al., 2025; van der Kuyl, 2026), is also observed here in the *N. coucang.* Nonetheless, similar to the most recently active ERV in humans (HERV-K), in *Nycticebus* genomes there appears to be ongoing replication of LERV2 and its subfamilies.

LERV2a in particular has a far greater number of CFL ERVs than solo LTRs, many of which have identical LTRs and ORFs. The *pol* and *env* LERV2a genes also show similar dN/dS values to an active retrovirus, HTLV, whilst the gag dN/dS value was higher than that observed in HTLV and higher than the LERV2a pol and env values. This may mean that an extracellular virus particle is unable to be formed through the function of *gag,* but the ERV is still able to replicate (e.g., by retrotransposition). The matrix domain of *gag,* with function in intracellular activity, shows higher purifying selection (lower dN/dS value), and the capsid domain, important for extracellular stability, has relaxed selection (Conte & Matthews, 1998; Kräusslich & Welker, 1996; Yamashita & Emerman, 2004). Through identifying unique TSDs for each LERV2 loci, thus showing unique insertions, we can reliably determine the recent insertion ages are not due to segmental duplication. Further evidence is supported by polymorphic insertions in the other *N. bengalensis* genome, where each genome contains shared and unique elements. Low dN/dS values do not necessarily reflect current or recent activity as exogenous retroviruses would be expected to be under strong purifying selection. Together, these findings are consistent with ongoing retrotransposition activity in LERV2a.

In other primates the most common genus of ERVs are betaretrovirus, type-D betaretrovirus and gammaretrovirus (Chabukswar et al., 2025; van der Kuyl, 2026) which is reflected in *N. coucang*. LERV2 and its closest relatives LERV3 and LERV4 belong to betaretroviruses as well as the previously discovered LERV1 (Michie et al., 2025). LERV2, LERV3 and LERV4 are most closely related to ERVs found in *Jaculus jaculus* and other rodents through BLASTs of all NCBI genomes. It is therefore likely that the ancestral virus originated from a rodent host, as rodents are well-established reservoirs for retroviruses for example KoRV (Mottaghinia et al., 2024; Simmons et al., 2014). Neither *J. jaculus* nor any of the other best BLASTn hits has a geographic range overlapping with that of slow lorises. Therefore, it is likely that a rodent species whose geographic range overlaps that of slow lorises has an ERV which is more closely related to LERV2, 3 and 4, and the lack of genomes from species in southeast Asia in general limits this analysis. It is also interesting to note that *J. jaculus* was the best hit in a similar study with PERVs (Chen et al., 2020), but the similarity was over 80%. LERV1 is shown again to be most closely related to HERV-K. The entirety of the HERV-K group has previously been reported to be absent in strepsirrhines and invasions occurred after the split of New World monkeys and Old World monkeys (∼40 MYA) (Barbulescu et al., 2001; Chabukswar et al., 2022). LERV1 is very distantly related to this HERV-K group however and only likely stems from a common ancestor with HERV-K.

Type-D betaretroviruses are well established cases of recombination where the retroviruses have a beta-type *pol* but a gamma-type *env* (Chabukswar et al., 2023; Henzy & Johnson, 2013). It is largely unknown how or why this type of recombination happens and is so common, but theories include expanding host range of the virus (Huder et al., 2002). Specifically, in primates it is thought that all primate type-D betaretroviruses originated from a single betaretroviral ancestor before the divergence of New World and Old World primates which gained a gammaretrovirus *env* from an unknown source leading to all known cases of type-D betaretroviruses in primates (Fine & Schochetman, 1978). Here five new families of type-D betaretroviruses were identified, three of which (LERV5, LERV6 and LERV8) are positioned with high confidence between the type-D betaretroviruses found in Old World primates (e.g., Simian retrovirus 2) and those found in New World primates (e.g., Squirrel monkey retrovirus). If the previous theory was true, the theory would likely need to be revised to encompass all primate lineages, especially given the presence of LERV7 which is more closely related to the New World than the old-world retroviruses. Evidence of recombination is not limited to phylogenetic discordance between families. In Loris tardigradus, all three LERV8 subfamilies retained >80% similarity in gag and pol relative to the N. coucang sequence, yet showed only ∼50% env similarity. This is consistent with env replacement in the L. tardigradus lineage following divergence from Nycticebus, further demonstrating that env recombination in type-D betaretroviruses can occur on relatively short evolutionary timescales within a single host lineage.

Sequences more closely related to LERV5 and LERV7 were identified through BLASTn searches against the NCBI mammalian genome database. LERV5 had closely related sequences to both rodents and Eulipotyphla, with *Erinaceus europaeus* having highest similarity. Both highest related species *(E. europeaus* and *Arvicathis niloticus*) do not have an overlapping range with slow lorises. LERV7 exhibits a broader phylogenetic distribution, with closely related sequences identified in both bats and *Manis javanica*, a species with a partially overlapping geographic range with slow lorises. Given the well-established role of bats as viral reservoirs, this pattern is consistent with a scenario in which a bat-associated retrovirus gave rise to related insertions in *M. javanica* and slow lorises. *M. javanica* has both an overlapping geographic range with slow lorises and is widely involved in the illegal wildlife trade (Challender et al., 2020; Nguyen et al., 2008; Zhang et al., 2015). However, cross-species transmission via modern wildlife trade is unlikely given the estimated insertion age of LERV7 (median = 10.48 MY). Instead, this pattern more likely suggests that an ancestral retrovirus infected both lineages historically, possibly through a shared vector species with overlapping ancestral ranges.

Amongst the gammaretroviruses there is an interesting relationship between LERV11 and LERV15 (hereafter referred to as LERV11/15), and their phylogenetic relationship. The relationship between LERV11 and LERV15 is complex as although both return similar BLASTn hits to bat and rodent genomes, these sequences are more similar to LERV15 than to LERV11, indicating asymmetric relatedness among the LERVs themselves. This suggests that LERV11 and LERV15 may derive from distinct but related viral lineages, or that differential divergence following insertion has obscured their original relationships. LERV10 was most closely related to HERV-E in the phylogenetic analysis. HERV-E itself has not been found in Strepsirrhini (Yi & Kim, 2006), it may be that a HERV-E related retrovirus infected the slow loris at a later date reflecting the estimated date of insertion (median = 23.42 MY) from a Haplorhini living in the area.

Sliding window analysis revealed an interesting trend where even in closest related LERVs (e.g. LERV2, LERV3 and LERV4) there is a large decrease in similarity in the region of the receptor binding domain (RBD) in *env*. A similar phenomenon has been observed in KoRV (Chappell et al., 2017; Greenwood et al., 2017), where they have different receptors to circumvent the hosts defence systems against previously infectious retroviruses. Alternatively, it may be that they are targeting different host cells. However, because LERV2, LERV3 and LERV4 share a common ancestor and LERV2 appears to have evolved directly from LERV3 based on the phylogenetic analysis, it is more likely the RBD region mutated to evade host defence systems.

LTRs were the most common differentiator between LERV subfamilies, where the internal genes of two subfamilies displayed over 80% identity, but the LTRs were more distinct. In humans and other primates approximately 90% of ERV loci are represented by solo LTRs (Buttler & Chuong, 2022; Y. Li et al., 2025) – this is not observed in the all LERVs we identify here. Traditionally, solo LTRs were thought to be 10-100 times more common than their full-length ERV counterparts (Bannert & Kurth, 2006), varying greatly among ERV families (Gemmell et al., 2019). Here we show a similar pattern of LERV diversity among the various families identified in the Nycticebus genome, as is observed amongst the larger primates.

This study presents the first comprehensive ERV catalogue for a strepsirrhine primate, revealing an unexpectedly diverse retroviral landscape in *Nycticebus* comprising 15 families and 34 subfamilies spanning three retroviral genera. Several findings have broad implications beyond the slow loris specifically. The identification of five novel type-D betaretrovirus families in a strepsirrhine lineage substantially extends the known diversity of this unusual recombinant group in primates, and the phylogenetic placement of LERV5, LERV6, and LERV8 between Old World and New World primate type-D betaretroviruses, combined with the distinct positioning of LERV7, collectively suggest that the evolutionary history of this group is more complex and more ancient than previously recognised. Most notably, evidence from LTR identity, dN/dS analysis, and insertional polymorphism between conspecific genomes is consistent with ongoing retrotransposition of LERV2a, demonstrating for the first time that potentially active ERV lineages are not exclusive to haplorrhines. This has implications for slow loris conservation and biosecurity: the frequent contact between slow lorises and humans through the illegal wildlife trade represents a potential route for cross-species retroviral exposure if endogenous elements were to give rise to infectious particles. Transcriptomic and proteomic validation of LERV2a activity, alongside broader genomic sampling across Southeast Asian mammal taxa, represent critical next steps. More broadly, this work demonstrates that the patterns of ERV accumulation, potentially active LERV families, and genomic distribution documented in haplorrhines are broadly conserved across the primate order, while also highlighting a distinct retroviral complement and ancestry that warrant further investigation across the suborder Strepsirrhini.

## 4. Methods

### 4.1 Identification of candidate ERVs by RepeatModeler

To identify repeat elements, RepeatModeler2 (RM2; Flynn et al., 2020) was run on the *N. coucang* reference genome (assembly accession GCA_027406575.1) using default parameters, with the long terminal repeat (LTR) structural discovery module enabled. Preliminary inspection of the output indicated that iterations 1–5 yielded few, if any, credible endogenous retrovirus (ERV) candidates and were therefore excluded from subsequent analyses. For example, in the 62 sequences that were produced from round 1, the mean length of the LTR was only 155bp, and through alignment with LTRs from the LTR Structural detection round most were only internal segments of LTRs.

Consensus sequences generated during the LTR discovery round were curated using a custom analysis pipeline (Figure 7). RM2 classifies consensus sequences as either Type=LTR, representing LTR-containing elements, or Type=INT, representing internal regions. In some cases, these classifications correspond to different components of the same ERV locus. To reduce redundancy, Type=LTR consensus sequences were matched to corresponding Type=INT consensus sequences, when present, based on the genomic coordinates of individual family members. When a Type=LTR consensus sequence could be associated with a corresponding Type=INT consensus sequence, only the Type=LTR consensus was retained. Consensus sequences classified as Type=LTR or Type=INT that lacked a corresponding match were retained independently for downstream analyses.

The retained Type=LTR and Type=INT consensus sequences were subsequently aligned to the corresponding reference genome using BLASTn. Searches were performed with an E-value threshold of 1 × 10LL and a minimum query coverage of 60%. To further reduce redundancy, BLASTn hits with genomic coordinates located within 10 bp of one another were considered to represent the same insertion, and only the hit with the highest percentage sequence identity was retained. To facilitate identification of potentially full-length ERVs, 14,000 bp of flanking sequence were extracted both upstream and downstream of each retained BLASTn hit. In some cases, RepeatModeler2 incorrectly classified consensus sequences as Type=LTR and Type=INT, for example, family 3 was annotated as Type=LTR despite representing an internal region based on the provided consensus sequence containing conserved domains of all genes. In such instances, the corresponding LTR consensus sequence was identified separately and used as an independent BLASTn query against the reference genome.

### 4.2 Candidate ERV annotation

NCBI’s conserved domain database (https://ftp.ncbi.nlm.nih.gov/pub/mmdb/cdd/little_endian/Cdd_NCBI_LE.tar.gz) was used (RPSTBLASTn) to identify the conserved domains within the genes of the potential ERV loci identified in the previous step (E-value cutoff of 1e^−5^). Open reading frames (ORFs) were identified in each potential ERV sequence using Geneious Prime V2025.2.2 (Kearse et al., 2012) with “Find ORF”, using a minimal length criterion of 399 nucleotides. To annotate LTRs within RM2-defined ERV families, a representative sequence from each family was aligned to itself with BLASTn to identify repeated terminal regions corresponding to putative LTRs.

Classification of loci and ERV families and subfamilies classification Sequences were classified into four categories: complete full-length (CFL), incomplete full-length, truncated, and solo LTR elements. A sequence was classified as CFL if it contained flanking 5′ and 3′ LTRs and at least one conserved domain corresponding to each of the major retroviral genes (*gag*, *pro*, *pol*, and *env*). Sequences that retained both flanking LTRs but lacked the expected conserved domains for one or more of these genes were classified as incomplete full-length. Sequences with some conserved domains but were lacking one LTR were classified as truncated elements. Sequences consisting of a single LTR and lacking any detectable retroviral conserved domains were classified as solo LTRs. Target site duplications (TSDs) were identified for each insertion, allowing for a maximum of one nucleotide mismatch between the duplicated sequences.

To identify subfamilies within each RM2-defined family, all complete full-length elements were aligned using MAFFT (Katoh and Standley, 2013). Alignments were manually inspected and refined, and alignment columns containing gaps in more than 95% of sequences were masked. Sequences containing insertions of other transposable elements, such as LINEs or unrelated LTR retrotransposons, were excluded from further analysis. Subfamily classification was based on separate nucleotide alignments of the LTR, gag, pro, pol, and env regions. Elements were assigned to distinct subfamilies partly following the 80-80-80 rule (Wicker et al., 2007) and criteria previously applied to Human endogenous retrovirus K (Subramanian et al., 2011). Initial clustering was performed using CD-HIT (Li and Godzik, 2006; Fu et al., 2012), followed by manual inspection. Consensus sequences were generated from aligning all CFL sequences for each family and subfamily and aligned to assess higher-level relationships among RM2 families. This comparison revealed that some RM2 families represented either the same biological family or subfamilies within a single family. Sequences were assigned to the same subfamily when the entire ERV (LTR, *gag, pro, pol* and *env*) had over 80% identity, and the same family when one or more regions had below 80% identity. Final families were called LERV (Loris endogenous retrovirus), followed by a number to represent family, followed by a letter to represent subfamily.

### 4.3 Population genetic analysis

#### 4.3.1 Estimation of insertion ages

The insertion age of each element was estimated based on sequence divergence between the 5′ and 3′ LTRs of complete full-length insertions. Insertion time was calculated using the formula: T=D/(2*r), where *D* represents LTR divergence (defined as the number of nucleotide differences divided by LTR length), and *r* is the neutral substitution rate of 1.72 × 10 substitutions per site per year estimated for lemurs (Campbell et al., 2021); there is no estimation for slow lorises or more closely related species. Large insertions and deletions (>10 bp) were excluded from LTR alignments prior to divergence estimation, as indels are not appropriately modelled under the neutral substitution framework underlying this equation.

#### 4.3.2 Phylogenetic analysis

Phylogenetic relationships of consensus full-length ERVs were inferred for each family and subfamily using alignments of conserved retroviral domains. Amino acid sequences corresponding to the reverse transcriptase (Pol), and transmembrane subunit (Env) domains were used for phylogenetic reconstruction of pro, pol, and env, respectively. Each dataset was aligned with representative retroviral sequences obtained from public databases (Supplementary Table 1). The reference dataset was enriched for betaretrovirus and gammaretrovirus sequences, as preliminary BLAST searches indicated that these genera were the most frequently recovered matches.

The best-fitting models of sequence evolution were selected using ModelFinder (Kalyaanamoorthy et al., 2017) implemented in IQ-TREE (Wong et al., 2025). Selected models were: gag – GTR+I+G4, Pro – RTREV+I+G4, Pol – RTREV+G5, and Env – LG+I+G4. Bayesian phylogenetic inference was performed using MrBayes (Huelsenbeck and Ronquist, 2001), with a burn-in of 100,000 generations and 1,000,000 Markov chain Monte Carlo (MCMC) iterations, using four chains (one cold and three heated) and a temperature parameter of 0.2. Trees were sampled every 200 generations. Effective sample sizes (ESS) of 832, 2206, 1735, and 999 for gag, pro, pol, and env, respectively, indicated convergence of the analyses. Resulting trees were visualised using TreeViewer (Bianchini and Sánchez-Baracaldo, 2024) and midpoint-rooted for presentation.

Following initial genus-level classification, additional phylogenetic analyses were performed within each major retroviral genus to resolve finer-scale relationships among endogenous and exogenous retroviruses (Supplementary Table 1). For Betaretrovirus and Gammaretrovirus, nucleotide-level alignments were used instead of amino acid sequences due to improved resolution at lower taxonomic levels. Additionally in few cases amino acid sequences were not accurately reconstructed due to indels within sequences. The same phylogenetic framework was applied, with ModelFinder identifying best-fitting substitution models (Betaretrovirus: gag – GTR+G4, pro – GTR+G4, pol – GTR+G5, env – GTR+I+G4; Gammaretrovirus: pol – GTR+I+G4). ESS values of 805, 1270, 727, and 1623 (Betaretrovirus) and 853, 842, 882, and 1057 (Gammaretrovirus) indicated convergence of Bayesian runs. All genus-level trees were rooted using Avian Leukemia Virus (KU375453) as an outgroup.

#### 4.3.3 Calculation of dN/dS ratios

The ratio of nonsynonymous (amino acid–altering) to synonymous (silent) substitutions (dN/dS) was used as an indicator of selective constraint, with lower values generally reflecting stronger purifying selection. As reliable dN/dS estimation requires complete and correctly annotated ORFs, only ERV families containing more than 10 full-length elements were included in this analysis to ensure robust reconstruction of coding sequences. Families in which the start or stop positions of any ORF could not be confidently determined were excluded. This filtering step resulted in six of the 15 identified families being retained for analysis, along with the LERV2 subfamilies.

To reconstruct intact coding sequences, point mutations affecting start codons or introducing premature stop codons were replaced with ambiguous nucleotides (“N”) to preserve reading frame continuity during translation. dN/dS ratios were estimated separately for each internal gene (*gag*, *pro*, *pol*, and *env*) using codon-based maximum likelihood methods implemented in CODEML (Yang, 2007). Due to frequent occurrence of zero-length branches in the reconstructed phylogenies, more complex branch models could not be reliably fitted. Consequently, only the one-ratio (M0) model was applied, providing a single dN/dS estimate averaged across all branches and sequences within each dataset. To further find evidence if any families were active, FEL (Murrell et al., 2012) was used to identify the proportion of codon sites that were under purifying selection.

#### 4.3.4 Identification of LERVs in related species

Consensus LERV sequences were used as queries in BLASTn searches against the genomes of progressively more distantly related species relative to *N. coucang*, until no further LERV sequences could be detected. A similarity threshold of 80% was applied to ensure that only homologous LERV insertions were retained, and a minimum query coverage of 90% was required to confirm that alignments spanned a substantial portion of the element, including the LTR regions, which are the primary basis for subfamily classification.

In total, six primate genomes were screened in a stepwise manner: *Nycticebus bengalensis* (GCA_023898255.1), *Xanthonycticebus pygmaeus* (GWHBCHX00000000), *Loris tardigradus* (GCA_023783135.1), *Perodicticus potto* (GCA_963574655.1), and *Otolemur garnettii* (GCA_000181295.3), with the search ultimately extending to *Lemur catta* (GCA_020740605.1), in which no LERV insertions were detected.

### 4.4 Identification of related ERVs in other species

Phylogenetic analyses provide a broad framework for assigning LERVs to established retroviral genera, but their interpretive power is limited by the availability of comparable ERV in public databases. As many mammalian genomes have not been systematically screened for ERVs, and annotated ERV sequences are often unavailable, additional similarity-based analyses were performed to identify the closest genomic relatives of selected LERV families.

This analysis focused on LERV families that were absent from *Loris tardigradus*, as these elements are more likely to represent relatively recent acquisitions in the *Nycticebus* lineage and therefore easier to find in other mammalian species. Full-length consensus sequences for each these LERV families were used as queries in BLASTn searches against the mammalian subset of the NCBI. To reduce taxonomic redundancy, only hits with the lowest E-value from each species were retained. The complete genomic region corresponding to each retained hit was extracted to allow comparison with putative full-length ERV insertions.

Extracted sequences were aligned with their corresponding LERV consensus sequences and analysed using SimPlot++ (Samson et al., 2022). Sliding-window similarity plots were generated to visualise variation in nucleotide identity across the length of each provirus. The hamming distance model with a sliding window size of 200bp and step size of 20bp was used to calculate similarity. This approach was used to determine whether genomic matches identified in other mammalian species showed consistently high similarity across the entire element or whether similarity was restricted to specific regions, such as pro and pol.

## Acknowledgments

C.A.G.M was funded by the Nigel Groome Studentship, Oxford Brookes University.

## References

Apakupakul, K., Deem, S. L., Maqsood, R., Sithiyopasakul, P., Wang, D., & Lim, E. S. (2021). Endogenization of a Prosimian Retrovirus during Lemur Evolution. Viruses, 13(3). 10.3390/v13030383

Bannert, N., & Kurth, R. (2006). The Evolutionary Dynamics of Human Endogenous Retroviral Families. Annual Review of Genomics and Human Genetics, 7(Volume 7, 2006), 149–173. 10.1146/annurev.genom.7.080505.115700

Barbulescu, M., Turner, G., Su, M., Kim, R., Jensen-Seaman, M. I., Deinard, A. S., Kidd, K. K., & Lenz, J. (2001). A HERV-K provirus in chimpanzees, bonobos and gorillas, but not humans. Current Biology: CB, 11(10), 779–783. 10.1016/s0960-9822(01)00227-5

Belshaw, R., Pereira, V., Katzourakis, A., Talbot, G., Pačes, J., Burt, A., & Tristem, M. (2004). Long-term reinfection of the human genome by endogenous retroviruses. Proceedings of the National Academy of Sciences, 101(14), 4894–4899. 10.1073/pnas.0307800101

Bénit, L., Lallemand, J.-B., Casella, J.-F., Philippe, H., & Heidmann, T. (1999). ERV-L Elements: A Family of Endogenous Retrovirus-Like Elements Active throughout the Evolution of Mammals. Journal of Virology, 73(4), 3301–3308. 10.1128/jvi.73.4.3301-3308.1999

Best, S., Le Tissier, P. R., & Stoye, J. P. (1997). Endogenous retroviruses and the evolution of resistance to retroviral infection. Trends in Microbiology, 5(8), 313–318. 10.1016/S0966-842X(97)01086-X

Blair, M. E., Cao, G. T. H., López-Nandam, E. H., Veronese-Paniagua, D. A., Birchette, M. G., Kenyon, M., Md-Zain, B. M., Munds, R. A., Nekaris, K. A.-I., Nijman, V., Roos, C., Thach, H. M., Sterling, E. J., & Le, M. D. (2023). Molecular Phylogenetic Relationships and Unveiling Novel Genetic Diversity among Slow and Pygmy Lorises, including Resurrection of Xanthonycticebus intermedius. Genes, 14(3), Article 3. 10.3390/genes14030643

Buttler, C. A., & Chuong, E. B. (2022). Emerging roles for endogenous retroviruses in immune epigenetic regulation. Immunological Reviews, 305(1), 165–178. 10.1111/imr.13042

Chabukswar, S., Grandi, N., Lin, L.-T., & Tramontano, E. (2023). Envelope Recombination: A Major Driver in Shaping Retroviral Diversification and Evolution within the Host Genome. Viruses, 15(9), 1856. 10.3390/v15091856

Chabukswar, S., Grandi, N., Soddu, E., Lin, L.-T., & Tramontano, E. (2025). Screening Envelope Genes Across Primate Genomes Reveals Evolution and Diversity Patterns of Endogenous Retroviruses. eLife, 13. 10.7554/eLife.104311.2

Chabukswar, S., Grandi, N., & Tramontano, E. (2022). Prolonged activity of HERV-K(HML2) in Old World Monkeys accounts for recent integrations and novel recombinant variants. Frontiers in Microbiology, 13. 10.3389/fmicb.2022.1040792

Challender, D. W. S., Heinrich, S., Shepherd, C. r., & Katsis, L. K. D. (2020). International trade and trafficking in pangolins, 1900–2019. In Pangolins (pp. 259–276). Academic Press. 10.1016/B978-0-12-815507-3.00016-2

Chappell, K. J., Brealey, J. C., Amarilla, A. A., Watterson, D., Hulse, L., Palmieri, C., Johnston, S. D., Holmes, E. C., Meers, J., & Young, P. R. (2017). Phylogenetic Diversity of Koala Retrovirus within a Wild Koala Population. Journal of Virology, 91(3), e01820–16. 10.1128/JVI.01820-16

Chen, Y., Chen, M., Duan, X., & Cui, J. (2020). Ancient origin and complex evolution of porcine endogenous retroviruses. Biosafety and Health, 02(03), 142–151. 10.1016/j.bsheal.2020.03.003

Chu, L., Su, F., Han, G.-Z., & Wang, J. (2023). Jawless vertebrates do not escape retrovirus infection. Virology, 583, 52–55. 10.1016/j.virol.2023.04.010

Chuong, E. B., Elde, N. C., & Feschotte, C. (2016). Regulatory evolution of innate immunity through co-option of endogenous retroviruses. Science, 351(6277), 1083–1087. 10.1126/science.aad5497

Conley, A. B., Piriyapongsa, J., & Jordan, I. K. (2008). Retroviral promoters in the human genome. Bioinformatics, 24(14), 1563–1567. 10.1093/bioinformatics/btn243

Conte, M. R., & Matthews, S. (1998). Retroviral Matrix Proteins: A Structural Perspective. Virology, 246(2), 191–198. 10.1006/viro.1998.9206

Fine, D., & Schochetman, G. (1978). Type D Primate Retroviruses: A Review1. Cancer Research, 38(10), 3123–3139.

Flynn, J. M., Hubley, R., Goubert, C., Rosen, J., Clark, A. G., Feschotte, C., & Smit, A. F. (2020). RepeatModeler2 for automated genomic discovery of transposable element families. Proceedings of the National Academy of Sciences, 117(17), 9451–9457. 10.1073/pnas.1921046117

Fu, L., Niu, B., Zhu, Z., Wu, S., & Li, W. (2012). CD-HIT: Accelerated for clustering the next-generation sequencing data. Bioinformatics, 28(23), 3150–3152. 10.1093/bioinformatics/bts565

Fuentes, D. R., Swigut, T., & Wysocka, J. (2018). Systematic perturbation of retroviral LTRs reveals widespread long-range effects on human gene regulation. eLife, 7, e35989. 10.7554/eLife.35989

Gemmell, P., Hein, J., & Katzourakis, A. (2019). The Exaptation of HERV-H: Evolutionary Analyses Reveal the Genomic Features of Highly Transcribed Elements. Frontiers in Immunology, 10. 10.3389/fimmu.2019.01339

Gifford, R. J., Blomberg, J., Coffin, J. M., Fan, H., Heidmann, T., Mayer, J., Stoye, J., Tristem, M., & Johnson, W. E. (2018). Nomenclature for endogenous retrovirus (ERV) loci. Retrovirology, 15(1), 59. 10.1186/s12977-018-0442-1

Gifford, R. J., Kabat, P., Martin, J., Lynch, C., & Tristem, M. (2005). Evolution and Distribution of Class II-Related Endogenous Retroviruses. Journal of Virology, 79(10), 6478–6486. 10.1128/jvi.79.10.6478-6486.2005

Gilbert, C., Maxfield, D. G., Goodman, S. M., & Feschotte, C. (2009). Parallel Germline Infiltration of a Lentivirus in Two Malagasy Lemurs. PLOS Genetics, 5(3), e1000425. 10.1371/journal.pgen.1000425

Greenwood, A. D., Ishida, Y., O’Brien, S. P., Roca, A. L., & Eiden, M. V. (2017). Transmission, Evolution, and Endogenization: Lessons Learned from Recent Retroviral Invasions. Microbiology and Molecular Biology Reviewsl : MMBR, 82(1), e00044–17. 10.1128/MMBR.00044-17

Groves, C. (1971). Systematics of the genus Nycticebus. Proceedings of the 3rd International Congress of Primatology, Zurich, 1971, 1, 44–53. https://cir.nii.ac.jp/crid/1571698600205759360

Groves, C. (2001). Primate Taxonomy. In Primate taxonomy (p. 99). Smithsonian Institution Press. http://bvbr.bib-bvb.de:8991/F?func=service&doc_library=BVB01&local_base=BVB01&doc_number=009619240&line_number=0001&func_code=DB_RECORDS&service_type=MEDIA

Henzy, J. E., & Johnson, W. E. (2013). Pushing the endogenous envelope. Philosophical Transactions of the Royal Society B: Biological Sciences, 368(1626), 20120506. 10.1098/rstb.2012.0506

Herniou, E., Martin, J., Miller, K., Cook, J., Wilkinson, M., & Tristem, M. (1998). Retroviral Diversity and Distribution in Vertebrates. Journal of Virology, 72(7), 5955–5966. 10.1128/jvi.72.7.5955-5966.1998

Holloway, J. R., Williams, Z. H., Freeman, M. M., Bulow, U., & Coffin, J. M. (2019). Gorillas have been infected with the HERV-K (HML-2) endogenous retrovirus much more recently than humans and chimpanzees. Proceedings of the National Academy of Sciences of the United States of America, 116(4), 1337–1346. 10.1073/pnas.1814203116

Huder, J. B., Böni, J., Hatt, J.-M., Soldati, G., Lutz, H., & Schüpbach, J. (2002). Identification and Characterization of Two Closely Related Unclassifiable Endogenous Retroviruses in Pythons (Python molurus and Python curtus). Journal of Virology, 76(15), 7607–7615. 10.1128/jvi.76.15.7607-7615.2002

Huelsenbeck, J. P., & Ronquist, F. (2001). MRBAYES: Bayesian inference of phylogenetic trees. Bioinformatics, 17(8), 754–755. 10.1093/bioinformatics/17.8.754

Izsvák, Z., Ma, J., Singh, M., & Hurst, L. (2025). Co-option of an endogenous retrovirus (LTR7- HERVH) in early human embryogenesis: Becoming useful and going unnoticed. Mobile DNA, 16. 10.1186/s13100-025-00361-0

Jern, P., & Coffin, J. M. (2008). Effects of Retroviruses on Host Genome Function. Annual Review of Genetics, 42(Volume 42, 2008), 709–732. 10.1146/annurev.genet.42.110807.091501

Kanda, R. K., Tristem, M., & Coulson, T. (2013). Exploring the effects of immunity and life history on the dynamics of an endogenous retrovirus. Philosophical Transactions of the Royal Society B: Biological Sciences, 368(1626), 20120505. 10.1098/rstb.2012.0505

Kassiotis, G., & Stoye, J. P. (2016). Immune responses to endogenous retroelements: Taking the bad with the good. Nature Reviews Immunology, 16(4), 207–219. 10.1038/nri.2016.27

Kassiotis, G., & Stoye, J. P. (2025). Evolution of antiviral host defenses against a backdrop of endogenous retroelements. Science, 389(6760), 588–593. 10.1126/science.adx4379

Katzourakis, A., & Gifford, R. J. (2010). Endogenous Viral Elements in Animal Genomes. PLOS Genetics, 6(11), e1001191. 10.1371/journal.pgen.1001191

Katzourakis, A., & Tristem, M. (2005). Phylogeny of human endogenous and exogenous retroviruses. In Retroviruses and Primate Genome Evolution (pp. 186–203). CRC Press.

Kearse, M., Moir, R., Wilson, A., Stones-Havas, S., Cheung, M., Sturrock, S., Buxton, S., Cooper, A., Markowitz, S., Duran, C., Thierer, T., Ashton, B., Meintjes, P., & Drummond, A. (2012). Geneious Basic: An integrated and extendable desktop software platform for the organization and analysis of sequence data. Bioinformatics, 28(12), 1647–1649. 10.1093/bioinformatics/bts199

Kessler, S. E., Tsangaras, K., Rasoloharijaona, S., Radespiel, U., & Greenwood, A. D. (2023). Long-term host–pathogen evolution of endogenous beta- and gammaretroviruses in mouse lemurs with little evidence of recent retroviral introgression. Virus Evolution, 9(1), veac117. 10.1093/ve/veac117

Kitao, K., Kryukov, K., Jin, L., Shintaku, Y., Hayakawa, T., & Nakagawa, S. (2025). Dynamic evolution of retroviral envelope-derived sequences in primates. 10.51094/jxiv.2051

Kräusslich, H.-G., & Welker, R. (1996). Intracellular Transport of Retroviral Capsid Components. In H.-G. Kräusslich (Ed.), Morphogenesis and Maturation of Retroviruses (pp. 25–63). Springer. 10.1007/978-3-642-80145-7_2

Lander, E. S., Linton, L. M., Birren, B., Nusbaum, C., Zody, M. C., Baldwin, J., Devon, K., Dewar, K., Doyle, M., FitzHugh, W., Funke, R., Gage, D., Harris, K., Heaford, A., Howland, J., Kann, L., Lehoczky, J., LeVine, R., McEwan, P., … The Wellcome Trust: (2001). Initial sequencing and analysis of the human genome. Nature, 409(6822), 860–921. 10.1038/35057062

Lavialle, C., Cornelis, G., Dupressoir, A., Esnault, C., Heidmann, O., Vernochet, C., & Heidmann, T. (2013). Paleovirology of ‘syncytins’, retroviral env genes exapted for a role in placentation. Philosophical Transactions of the Royal Society B: Biological Sciences, 368(1626), 20120507. 10.1098/rstb.2012.0507

Lee, A., Huntley, D., Aiewsakun, P., Kanda, R. K., Lynn, C., & Tristem, M. (2014). Novel Denisovan and Neanderthal retroviruses. Journal of Virology, 88(21), 12907–12909. 10.1128/JVI.01825-14

Li, W., & Godzik, A. (2006). Cd-hit: A fast program for clustering and comparing large sets of protein or nucleotide sequences. Bioinformatics, 22(13), 1658–1659. 10.1093/bioinformatics/btl158

Li, Y., Jiang, C., Li, L., Zhang, Z., He, Y., He, K., Lu, Z., Huang, Y., & Liang, B. (2025). A Compendium of ERVs in the Genomes of Major Primates. iCell, 2(2). 10.71373/SBHJ1439

Li, Y., Zhang, G., & Cui, J. (2022). Origin and Deep Evolution of Human Endogenous Retroviruses in Pan-Primates. Viruses, 14(7), Article 7. 10.3390/v14071370

Mayer, J., Blomberg, J., & Seal, R. L. (2011). A revised nomenclature for transcribed human endogenous retroviral loci. Mobile DNA, 2(1), 7. 10.1186/1759-8753-2-7

McCarthy, E. M., & McDonald, J. F. (2004). Long terminal repeat retrotransposons of Mus musculus. Genome Biology, 5(3), R14. 10.1186/gb-2004-5-3-r14

Medstrand, P., & Mager, D. L. (1998). Human-Specific Integrations of the HERV-K Endogenous Retrovirus Family. Journal of Virology, 72(12), 9782–9787. 10.1128/jvi.72.12.9782-9787.1998

Mi, S., Lee, X., Li, X., Veldman, G. M., Finnerty, H., Racie, L., LaVallie, E., Tang, X.-Y., Edouard, P., Howes, S., Keith, J. C., & McCoy, J. M. (2000). Syncytin is a captive retroviral envelope protein involved in human placental morphogenesis. Nature, 403(6771), 785–789. 10.1038/35001608

Michie, C. A. G., Free, H. B., Nijman, V., & Kanda, R. K. (2025). A novel endogenous retrovirus in slow lorises (Nycticebus) and its role in species identification. Virology, 610, 110600. 10.1016/j.virol.2025.110600

Moore, R. S., Wihermanto, & Nekaris, K. a. I. (2014). Compassionate conservation, rehabilitation and translocation of Indonesian slow lorises. Endangered Species Research, 26(2), 93–102. 10.3354/esr00620

Mottaghinia, S., Stenzel, S., Tsangaras, K., Nikolaidis, N., Laue, M., Müller, K., Hölscher, H., Löber, U., McEwen, G. K., Donnellan, S. C., Rowe, K. C., Aplin, K. P., Goffinet, C., & Greenwood, A. D. (2024). A recent gibbon ape leukemia virus germline integration in a rodent from New Guinea. Proceedings of the National Academy of Sciences, 121(6), e2220392121. 10.1073/pnas.2220392121

Murrell, B., Wertheim, J. O., Moola, S., Weighill, T., Scheffler, K., & Pond, S. L. K. (2012). Detecting Individual Sites Subject to Episodic Diversifying Selection. PLOS Genetics, 8(7), e1002764. 10.1371/journal.pgen.1002764

Nekaris, K. A. I., & Burrows, A. M. (2020). Evolution, Ecology and Conservation of Lorises and Pottos’. Cambridge University Press.

Nekaris, K. A. I., Shepherd, C. r., Starr, C. r., & Nijman, V. (2010). Exploring cultural drivers for wildlife trade via an ethnoprimatological approach: A case study of slender and slow lorises (Loris and Nycticebus) in South and Southeast Asia. American Journal of Primatology, 72(10), 877–886. 10.1002/ajp.20842

Nekaris, K. A. I., & Starr, C. R. (2015). Conservation and ecology of the neglected slow loris: Priorities and prospects. Endangered Species Research, 28(1), 87–95. 10.3354/esr00674

Nguyen, T., Newton, P., Roberton, S., Wicker, L., & Bell, D. (2008). Tapping into local knowledge to help conserve pangolins in Vietnam.

Palmada-Flores, M., Orkin, J. D., Haase, B., Mountcastle, J., Bertelsen, M. F., Fedrigo, O., Kuderna, L. F. K., Jarvis, E. D., & Marques-Bonet, T. (2022). A high-quality, long-read genome assembly of the endangered ring-tailed lemur (Lemur catta). GigaScience, 11, giac026. 10.1093/gigascience/giac026

Patzke, S., Lindeskog, M., Munthe, E., & Aasheim, H.-C. (2002). Characterization of a Novel Human Endogenous Retrovirus, HERV-H/F, Expressed in Human Leukemia Cell Lines. Virology, 303(1), 164–173. 10.1006/viro.2002.1615

Pozzi, L., Nekaris, K. A.-I., Perkin, A., Bearder, S. K., Pimley, E. R., Schulze, H., Streicher, U., Nadler, T., Kitchener, A., Zischler, H., Zinner, D., & Roos, C. (2015). Remarkable ancient divergences amongst neglected lorisiform primates: Phylogeny of Lorisiform Primates. Zoological Journal of the Linnean Society, 175(3), 661–674. 10.1111/zoj.12286

Romano, C. M., de Melo, F. L., Corsini, M. A. B., Holmes, E. C., & Zanotto, P. M. de A. (2007). Demographic histories of ERV-K in humans, chimpanzees and rhesus monkeys. PloS One, 2(10), e1026. 10.1371/journal.pone.0001026

Samson, S., Lord, É., & Makarenkov, V. (2022). SimPlot++: A Python application for representing sequence similarity and detecting recombination. Bioinformatics, 38(11), 3118–3120. 10.1093/bioinformatics/btac287

Simmons, G., Clarke, D., McKee, J., Young, P., & Meers, J. (2014). Discovery of a Novel Retrovirus Sequence in an Australian Native Rodent (Melomys burtoni): A Putative Link between Gibbon Ape Leukemia Virus and Koala Retrovirus. PLOS ONE, 9(9), e106954. 10.1371/journal.pone.0106954

Starr, C., Nekaris, K. a. I., Streicher, U., & Leung, L. (2010). Traditional use of slow lorises Nycticebus bengalensis and N. pygmaeus in Cambodia: An impediment to their conservation. Endangered Species Research, 12, 17–23. 10.3354/esr00285

Subramanian, R. P., Wildschutte, J. H., Russo, C., & Coffin, J. M. (2011). Identification, characterization, and comparative genomic distribution of the HERV-K (HML-2) group of human endogenous retroviruses. Retrovirology, 8, 90. 10.1186/1742-4690-8-90

Thlllch, H. M., Le, M. D., Vũ, N. B., Panariello, A., Sethi, G., Sterling, E. J., & Blair, M. E. (2018). Slow Loris Trade in Vietnam: Exploring Diverse Knowledge and Values. Folia Primatologica, 89(1), 45–62. 10.1159/000481196

van der Kuyl, A. C. (2026). Gammaretrovirus Infections in Humans in the Past, Present, and Future: Have We Defeated the Pathogen? Pathogens, 15(1), 104. 10.3390/pathogens15010104

Wang, J., & Han, G.-Z. (2023). Genome mining shows that retroviruses are pervasively invading vertebrate genomes. Nature Communications, 14(1), 4968. 10.1038/s41467-023-40732-w

Wildschutte, J. H., Williams, Z. H., Montesion, M., Subramanian, R. P., Kidd, J. M., & Coffin, J. M. (2016). Discovery of unfixed endogenous retrovirus insertions in diverse human populations. Proceedings of the National Academy of Sciences, 113(16), E2326–E2334. 10.1073/pnas.1602336113

Yamashita, M., & Emerman, M. (2004). Capsid Is a Dominant Determinant of Retrovirus Infectivity in Nondividing Cells. Journal of Virology, 78(11), 5670–5678. 10.1128/jvi.78.11.5670-5678.2004

Yang, Z. (2007). PAML 4: Phylogenetic Analysis by Maximum Likelihood. Molecular Biology and Evolution, 24(8), 1586–1591. 10.1093/molbev/msm088

Yi, J.-M., & Kim, H.-S. (2006). Molecular evolution of the HERV-E family in primates. Archives of Virology, 151(6), 1107–1116. 10.1007/s00705-005-0701-z

Zhang, H., Miller, M. P., Yang, F., Chan, H. K., Gaubert, P., Ades, G., & Fischer, G. A. (2015). Molecular tracing of confiscated pangolin scales for conservation and illegal trade monitoring in Southeast Asia. Global Ecology and Conservation, 4, 414–422. 10.1016/j.gecco.2015.08.002

Zheng, W., Gojobori, J., Suh, A., & Satta, Y. (2024). Different Host–Endogenous Retrovirus Relationships between Mammals and Birds Reflected in Genome-Wide Evolutionary Interaction Patterns. Genome Biology and Evolution, 16(4), evae065. 10.1093/gbe/evae065

