## Supplementary information for "Abundance, diversity and activity of endogenous retroviruses in the slow loris"

**Supplementary Table 1: Sequences used in phylogenetic analysis of LERV seqeunces. N/A indicates not included in analysis for the gene. What type of sequence was used and available is shown.**

| **Name** | **Accession number (NCBI or Dfam) or reference** | **Gag genera** | **Pro genera** | **Pol genera** | **Env genera** | **Nucleotide or amino acid tree** |
| --- | --- | --- | --- | --- | --- | --- |
| Avian leukaemia virus | NC_015116 | Alpha | Alpha | Alpha | Alpha | Amino acid |
| Avian leukosis virus | KU375453 | Alpha | Alpha | Alpha | Alpha | Amino acid and nucleotide outgroup |
| Black Syrian hamster retrovirus | MK304634 | Beta | Beta | Beta | Beta | Both |
| Enzootic nasal tumour virus 2 | MK164396 | Beta | Beta | Beta | Beta | Both |
| Gorilla HERV-K HML-2-12 | MH768776 | Beta | Beta | Beta | Beta | Nucleotide |
| Gorilla HERV-K HML-2-9 | MH678771 | Beta | Beta | Beta | Beta | Nucleotide |
| HERV-K HML-10 | (Grandi et al., 2019) | Beta | N/A | Beta | Beta | Both |
| HERV-K HML-2 | JN675029 | Beta | Beta | Beta | Beta | Both |
| HERV-K HML-5 | (Lavie et al., 2004) | Beta | Beta | Beta | Beta | Both |
| HERV-K HML-8 | DF000000189 | Beta | Beta | Beta | Beta | Both |
| HERV-K HML-9 | DF000000193 | Beta | Beta | Beta | Beta | Both |
| HERV-K113 | NC_022518 | Beta | Beta | Beta | Beta | Both |
| HERV-K115 | AY037929 | Beta | Beta | Beta | Beta | Both |
| Jaagsiekte sheep retrovirus | NC_001494 | Beta | Beta | Beta | Beta | Both |
| Marsupial endogenous betaretrovirus 3 | BK068961 | Beta | Beta | Beta | Beta | Nucleotide |
| Mouse mammary tumor virus | NC_001503 | Beta | Beta | Beta | Beta | Both |
| Ovine enzootic nasal tumour virus | NC_007015 | Beta | Beta | Beta | Beta | Both |
| Yunnan shrew betaretrovirus 1 | PV979745 | Beta | Beta | Beta | Beta | Nucleotide |
| Desmodus rotundus ERV | NC_027117 | Beta | Beta | Beta | Gamma | Both |
| Mason-Pfizer monkey virus | NC_001550 | Beta | Beta | Beta | Gamma | Both |
| Prosimian retrovirus 1 | MT787217 | Beta | Beta | Beta | Gamma | Both |
| Python molorus endogenous retrovirus | AF500296 | Beta | Beta | Beta | Gamma | Nucleotide |
| Simian endogenous retrovirus | U85505 | Beta | Beta | Beta | Gamma | Both |
| Simian retrovirus 1 | M11841 | Beta | Beta | Beta | Gamma | Both |
| Simian retrovirus 2 | MF805808 | Beta | Beta | Beta | Gamma | Both |
| Simian retrovirus 4 | NC_014474 | Beta | Beta | Beta | Gamma | Both |
| Simian retrovirus 8 | NC_031326 | Beta | Beta | Beta | Gamma | Both |
| Squirrel monkey retrovirus | NC_001514 | Beta | Beta | Beta | Gamma | Both |
| Cervid endogenous betaretrovirus 1 | OL547611.1 | Beta | Beta | Beta | N/A | Amino acid |
| HERV-K HML-1 | AH013884 | N/A | N/A | Beta | N/A | Both |
| HERV-K HML-3 | U35153 | N/A | N/A | Beta | N/A | Both |
| HERV-K HML-4 | AF020092 | Beta | Beta | Beta | N/A | Both |
| HERV-K HML-6 | (Medstrand et al., 1997) | Beta | N/A | Beta | N/A | Both |
| HERV-K HML-7 | AF016000 | N/A | N/A | Beta | N/A | Both |
| RV_Slow_Loris | (R. Gifford et al., 2005) | N/A | N/A | Beta | N/A | Amino acid |
| Human T-lymphotropic virus 2 | NC_001488 | Delta | Delta | Delta | Delta | Amino acid |
| Simian T-cell lymphotropic | NC_011546 | Delta | Delta | Delta | Delta | Amino acid |
| Avian reticuloendotheiosis virus | NC_006934 | Gamma | Gamma | Gamma | Gamma | Nucleotide |
| Baboon endogenous virus | NC_022517 | Gamma | Gamma | Gamma | Gamma | Both |
| ERV-Gamma-jja | Y. Chen et al., 2020 | Gamma | Gamma | Gamma | Gamma | Nucleotide |
| Feline Leukemia virus | NC_001940 | Gamma | Gamma | Gamma | Gamma | Both |
| Friend murine leukemia virus | NC_001362 | Gamma | Gamma | Gamma | Gamma | Nucleotide |
| Gibbon ape leukaemia virus | NC_001885 | Gamma | Gamma | Gamma | Gamma | Both |
| Grey mouse lemur ERV-Fc | (Diehl et al., 2016) | N/A | N/A | Gamma | Gamma | Amino acid |
| HERV-E | FM212573 | Gamma | Gamma | Gamma | Gamma | Nucleotide |
| HERV-H | AJ289709 | Gamma | Gamma | Gamma | Gamma | Nucleotide |
| HERV-T1 | AY836222 | Gamma | Gamma | Gamma | Gamma | Nucleotide |
| HERV-W | AY101582 | N/A | Gamma | Gamma | Gamma | Nucleotide |
| Koala retrovirus | NC_039228 | Gamma | Gamma | Gamma | Gamma | Both |
| Orcinus orca ERV | GQ22416 | Gamma | Gamma | Gamma | Gamma | Nucleotide |
| PERV-A | AJ293656 | Gamma | Gamma | Gamma | Gamma | Both |
| PERV-B | AY099324 | Gamma | Gamma | Gamma | Gamma | Nucleotide |
| PERV-C | AJ293656 | Gamma | Gamma | Gamma | Gamma | Nucleotide |
| Rauscher murine leukemia virus | NC_001819 | Gamma | Gamma | Gamma | Gamma | Nucleotide |
| Rhinolophus ferremequinum retrovirus | JQ303225 | Gamma | Gamma | Gamma | Gamma | Nucleotide |
| Woolly monkey sarcoma virus | KT724051 | Gamma | Gamma | Gamma | Gamma | Both |
| HERV-T2 | AY836223 | Gamma | Gamma | Gamma | N/A | Nucleotide |
| RD114 retrovirus | NC_009889 | Gamma | Gamma | Gamma | N/A | Nucleotide |
| Gorilla HERV-T | AB266802 | N/A | N/A | N/A | Gamma | Nucleotide |
| Mus dunni ERV | AF053745 | N/A | N/A | N/A | Gamma | Nucleotide |
| Brown greater galago prosimian foamy virus | NC_039023 | Spuma | Spuma | Spuma | N/A | Amino acid |
| Rhesus macaque simian foamy virus | NC_039238 | Spuma | Spuma | Spuma | N/A | Amino acid |

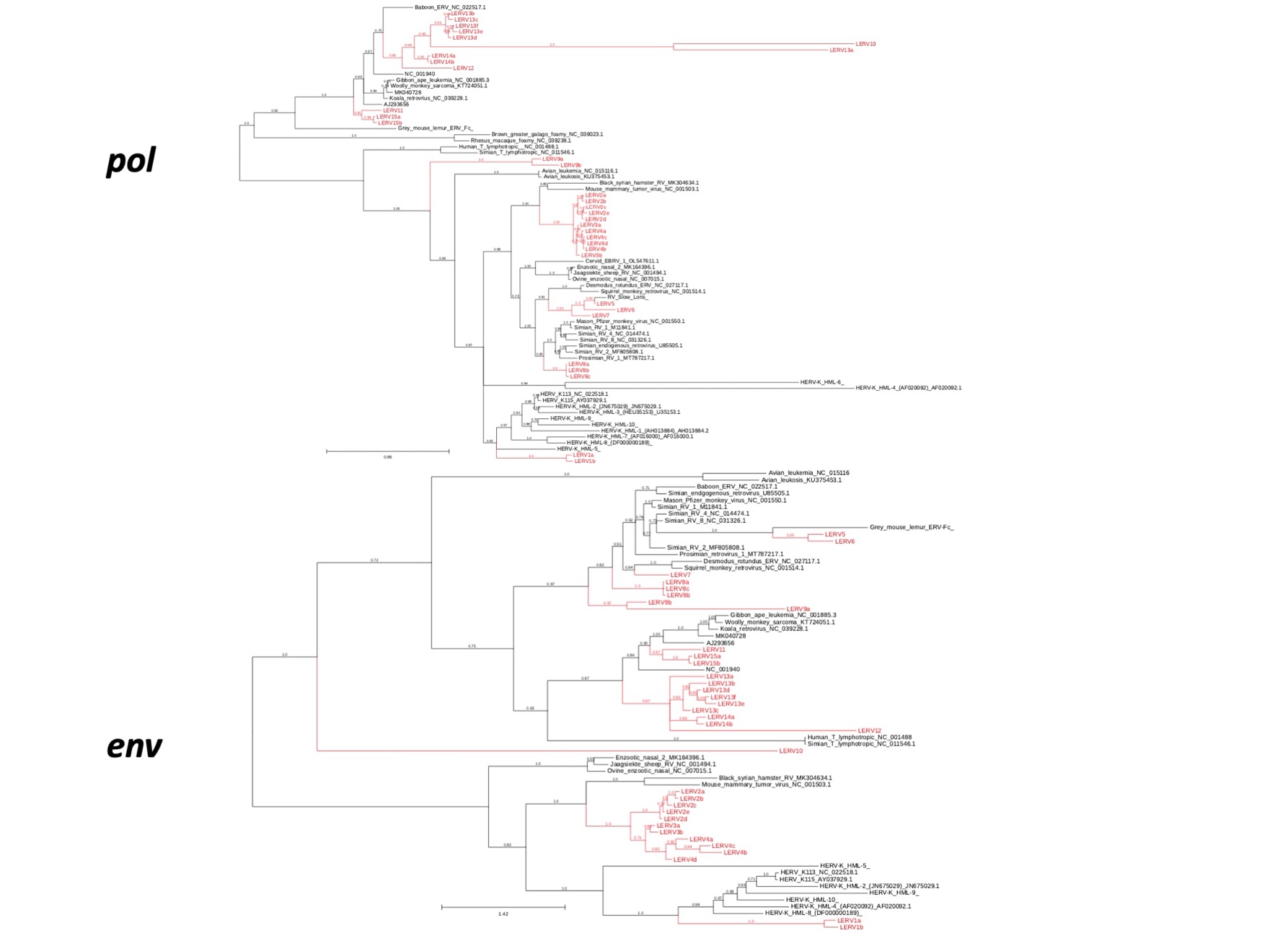

**Supplementary Figure 1: Bayesian phylogenetic tree of LERV families inferred from *pol* and *env* amino acid regions of reverse transcriptase and the transmembrane subunit. LERV families are highlighted in red. Trees have been rooted at midpoint.**

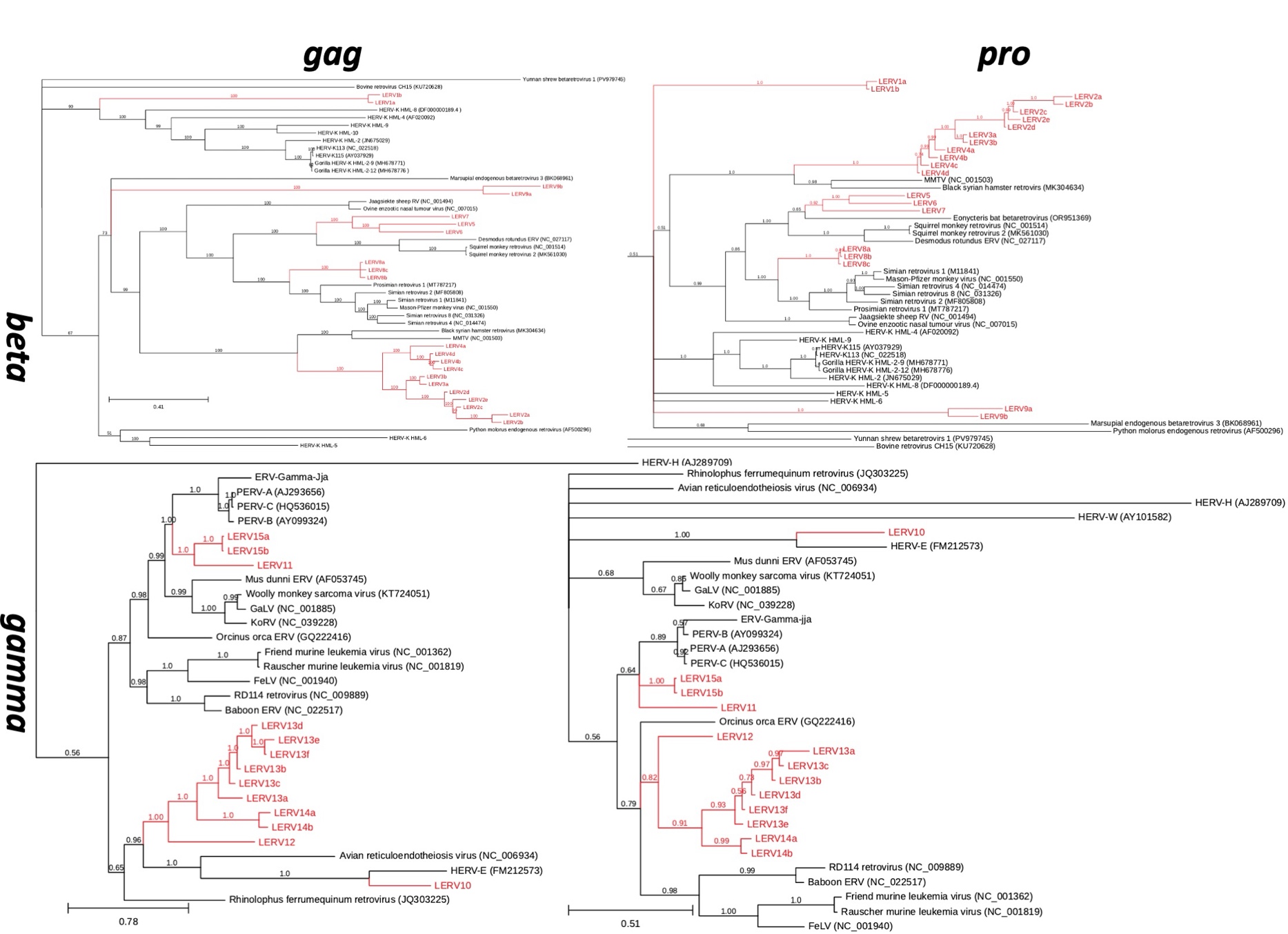

**Supplementary Figure 2: Bayesian phylogenetic tree of LERV families inferred from gag and pro whole nucleotide region rooted using Avian Leukosis Virus as an outgroup (not shown for visual purposes. LERV families are highlighted in red.**

**Supplementary table 2: Human T-cell lymophotropic virus sequences used in dN/dS analysis.**

**NCBI accession number**

LC192501

LC192533

LC192517

LC192518

LC192500

LC192506

LC192507

LC192532

LC192524

LC192522

LC192516

LC192521

LC192523

LC192535

LC192531

LC192504

LC192505

LC192525

LC192519

LC192502

LC192520

LC192503

LC192530

LC192510

LC192512

LC192534

LC192509

LC192508

LC192511

LC192536

LC192514

LC192528

LC192513

LC192526

LC192527

LC192515

LC192529

AF139170

MN781155

MN781153

KX430030

KX430031

MN781150

MN781149

MN781154

MN781156

KF242506

KX905203

KX905202

**Supplementary Table 3: Distribution of LERV insertions across chromosomes of the reference *N. coucang* genome. Enrichment value is the number of observed insertions divided by the number of expected insertions.**

| Chromosome | Chromosome Length | Observed LERV insertions | Expected number of insertions | Binomial p-value | BH adjusted p-value | Enrichment value |
| --- | --- | --- | --- | --- | --- | --- |
| 1 | 196249132 | 422 | 422.731 | 1.000 | 1.000 | 0.998 |
| 2 | 184820795 | 277 | 398.113 | 0.000 | 0.000 | 0.696 |
| 3 | 161203717 | 281 | 347.241 | 0.000 | 0.001 | 0.809 |
| 4 | 148538759 | 277 | 319.960 | 0.014 | 0.026 | 0.866 |
| 5 | 142425405 | 299 | 306.792 | 0.682 | 0.775 | 0.975 |
| 6 | 138330575 | 304 | 297.971 | 0.722 | 0.784 | 1.020 |
| 7 | 137715312 | 236 | 296.646 | 0.000 | 0.001 | 0.796 |
| 8 | 136588019 | 247 | 294.217 | 0.004 | 0.010 | 0.840 |
| 9 | 135917784 | 257 | 292.774 | 0.031 | 0.049 | 0.878 |
| 10 | 133885751 | 287 | 288.397 | 0.976 | 1.000 | 0.995 |
| 11 | 128238841 | 235 | 276.233 | 0.011 | 0.022 | 0.851 |
| 12 | 112708720 | 169 | 242.780 | 0.000 | 0.000 | 0.696 |
| 13 | 103192619 | 213 | 222.282 | 0.561 | 0.668 | 0.958 |
| 14 | 99988313 | 204 | 215.380 | 0.446 | 0.577 | 0.947 |
| 15 | 97885895 | 221 | 210.851 | 0.462 | 0.577 | 1.048 |
| 16 | 96499272 | 174 | 207.864 | 0.016 | 0.027 | 0.837 |
| 17 | 91048355 | 155 | 196.123 | 0.002 | 0.006 | 0.790 |
| 18 | 80579731 | 117 | 173.573 | 0.000 | 0.000 | 0.674 |
| 19 | 72490086 | 172 | 156.147 | 0.195 | 0.286 | 1.102 |
| 20 | 69287667 | 349 | 149.249 | 0.000 | 0.000 | 2.338 |
| 21 | 67625389 | 117 | 145.669 | 0.015 | 0.027 | 0.803 |
| 22 | 61455708 | 85 | 132.379 | 0.000 | 0.000 | 0.642 |
| 23 | 38949463 | 60 | 83.899 | 0.007 | 0.016 | 0.715 |
| 24 | 33060932 | 62 | 71.215 | 0.310 | 0.431 | 0.871 |
| X | 187328519 | 593 | 403.515 | 0.000 | 0.000 | 1.470 |
